# System-wide dissociation of reward and aversive dopaminergic signals

**DOI:** 10.1101/2025.05.27.656467

**Authors:** Sarah-Julie Bouchard, Joël Boutin, Martin Lévesque, Vincent Breton-Provencher

## Abstract

Dopamine is central to reinforcement learning, classically linked to reward prediction error signaling. However, recent findings suggest a more complex picture, with dopamine neurons responding to aversive or non-reward-related events and displaying diverse projection patterns. To investigate how dopamine release varies across the brain, we used multi-site fiber photometry to record the fluorescent dopamine sensor GRAB-DA in mice performing tasks involving both rewarding and aversive outcomes. We found that while reward-related dopamine signals were broadly distributed, responses to aversive events were region-specific, enabling classification of projection targets from these dopamine dynamics. By using dimensionality reduction to examine the main axis of covariance of dopamine release across the brain, we found that reward prediction error signals are encoded by parallel manifolds, whereas aversive stimulus signals are encoded by orthogonal manifolds. Thus, our findings support a distributed, vector-valued model of dopaminergic signaling, in which anatomically and functionally distinct pathways contribute to encoding of reinforcement-related variables across the brain.

## Introduction

The dopamine system has long been implicated in reinforcement learning, particularly through its role in signaling reward prediction error – transient increases or decreases in activity following outcomes that are better or worse than expected, respectively^1–3^. However, numerous studies have shown that a subset of dopamine neurons also respond to aversive stimuli^4–7^, or other non-reward related events^8,9^, challenging the notion of a unitary, scalar signal broadcasted to all dopaminergic outputs and instead suggesting a heterogeneous framework for dopamine function.

Anatomical and molecular evidence indicates that dopamine neurons are functionally diverse. Located primarily in the substantia nigra (SN) and ventral tegmental area (VTA), dopamine neurons project broadly to the striatum, cortex, and limbic areas^10^. Single-cell gene expression profiling has revealed transcriptionally distinct dopamine subtypes that, to a certain extent, correspond to specific projection targets^11,12^. Consistent with this, anatomical tracing studies show that dopamine axons projecting to the striatum form dense and spatially confined arbors with little collateralization^13–16^, while projections to the prefrontal cortex or amygdala are more diffuse and extensively collateralized^15^. Yet, input-mapping studies reveal largely overlapping afferents across projection-defined dopamine populations^17–19^, therefore, differential dopamine release in distinct brain areas cannot be predicted solely from anatomical evidence.

Recent studies of projection-defined dopamine axons have begun to clarify whether prediction error signals are conveyed by distinct dopaminergic pathways. Within the striatum, dopaminergic activity gradients reflect different reinforcement learning features: anterior and ventral regions show canonical reward prediction error signals^20–23^, while posterior areas display signals related to salience and novelty^23–25^. Medial and lateral striatal regions may also exhibit functional gradients, with prediction error signals localized more laterally^26–28^. Beyond the striatum, dopamine axons in the medial prefrontal cortex and amygdala encode reward-related signals and aversive signals^29–32^. These findings suggest that dopamine release varies by stimulus valence and reward expectation, but also with projection target. However, these studies often focus on single regions and use distinct behavioral paradigms, limiting our understanding of how dopaminergic signals relate across the striatum, limbic system, and cortex. Determining how dopamine release varies across targets is crucial for identifying unifying coding schemes for region-specific signals driving learning and behavior.

In this study, we sought to systematically map dopamine release across the major targets of the dopaminergic system in reinforcement learning. Using multi-site fiber photometry recordings of a fluorescent dopamine sensor, GRAB-DA^33^, we simultaneously measured the temporal dynamics of dopamine release across several brain regions in response to rewarding and aversive outcomes. We found that reward prediction error signals were relatively distributed, while aversive signals were more spatially selective. Dimensionality reduction revealed that dopamine signals to different reinforcers occupy distinct subspaces, suggesting that cross-region covariation may be a mechanism by which dopamine separates prediction error from other non-reward related features.

These results support the idea that dopamine conveys a distributed reinforcement signal that enables parallel learning processes across the brain.

## Results

### Multi-site recordings of fast dopamine dynamics

To record dopamine release in various brain locations, we expressed GRAB-DA3m^33^, and tracked temporal dynamics of dopamine release with multi-site, fiber photometry (**Fig. 1A**). For each recording, we aligned the position of the tip of the optic fiber to coronal sections of the Allen Reference Atlas^34^ using point registration (**Fig. 1B,C, S1A**). Using this approach, we targeted the dorsal striatum (DS), tail striatum (TS), and the lateral, core, and medial parts of the nucleus accumbens (NAc_lat_, NAc_c_, and NAc_m_, respectively), olfactory tubercle (OT), basolateral amygdala (BLA), and medial prefrontal cortex (mPFC) (**Fig. 1C**). This selection of targets allowed us to sample signals along the nigrostriatal (DS, TS), mesolimbic (NAc, OT), and mesocorticolimbic pathways (BLA, mPFC). Furthermore, each pathway was sampled to facilitate comparisons of dopamine release across regions with known molecular heterogeneity^11^.

**Figure 1.**
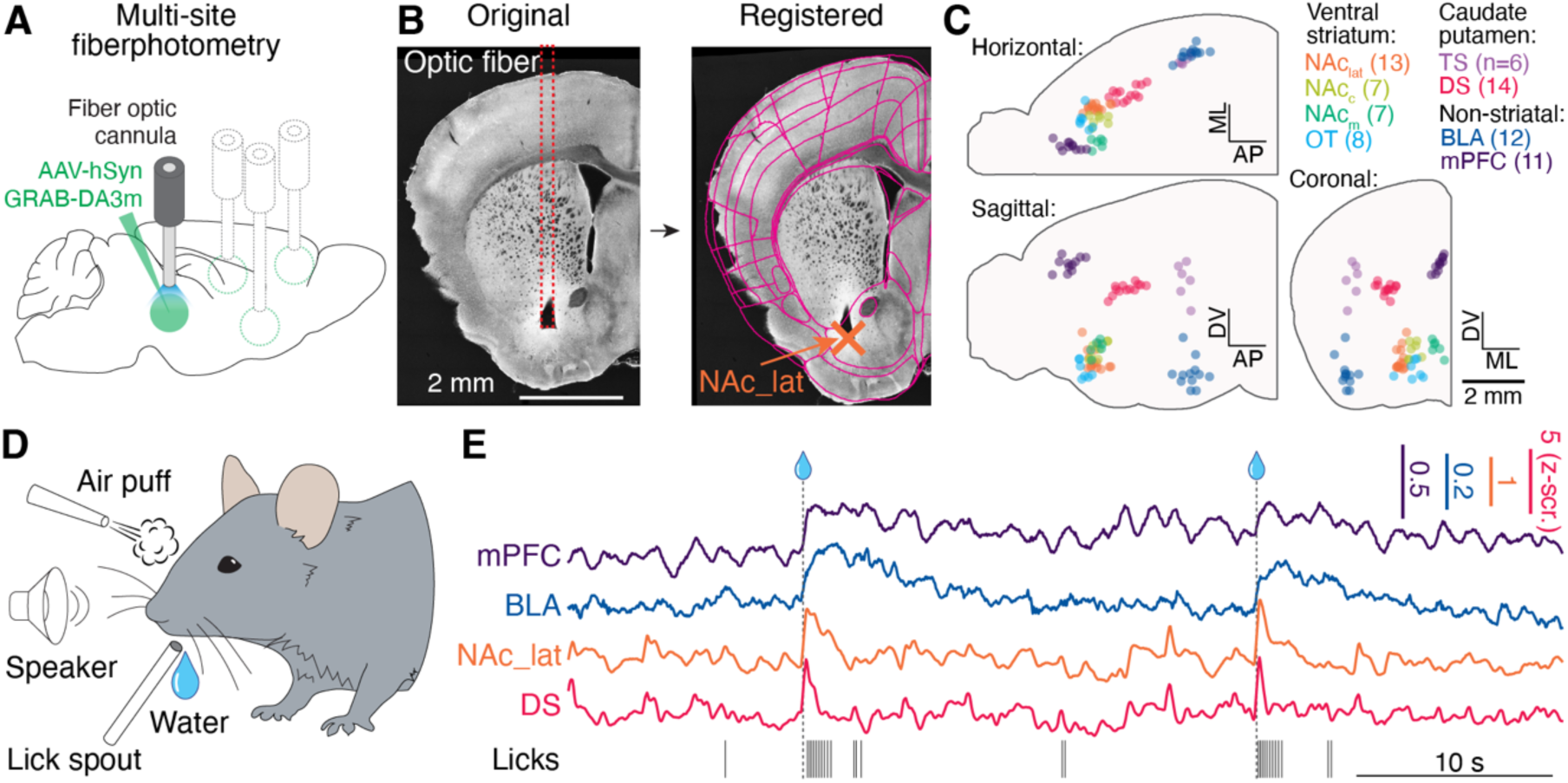
Multi-site recordings of dopamine signals. **(A)** Schematic of the surgical strategy for viral injection of GRAB-DA and fiber optic implantation. **(B)** Left: A representative coronal slice showing fiber tip locations (red dashed lines) in the NAc_lat_. Scale bar, 2 mm. Right: Slice alignment to the Allen Brain Atlas using point registration (atlas outlines in magenta). See also **Figure S1A. (C)** Anatomical locations of all fiber tips used in this study (N = 78 recording sites). Scale bar, 2 mm. **(D)** Schematic of the behavioral apparatus. **(E)** Representative GRAB-DA traces recorded simultaneously in mPFC, BLA, NAc_lat_, and DS. Vertical ticks at the bottom indicate individual licks; vertical dashed lines indicate time of water reward delivery.

Our fiber photometry system enabled us to simultaneously record dopamine in up to four brain regions in head-restrained mice performing various behavioral paradigms (**Figs. 1D,E, S1B**). To validate the ability of our system to reliably detect dopamine release in regions with sparse dopaminergic axons, such as the mPFC^35,36^, we combined optogenetics with fiber photometry. We used Dat-IRES-Cre mice to specifically express Chrmine2.0^37^, a red-shifted activating opsin, in dopaminergic neurons of either the VTA or the SN and implanted an optic fiber above the injection site (**Fig. S1C**). A second optic fiber was used to measure the GRAB-DA signal in a target area in response to photoactivation of dopaminergic neurons (**Fig. S1D**). A single 10-ms pulse of light evoked reliable GRAB-DA signals not only in the NAc_lat_ and DS, as expected, but also in the mPFC (**Fig. S1E**).

Next, we used photoinhibition to assess the ability of our experimental approach to detect dopamine over noradrenaline signals in regions where both neuromodulators are present, such as the mPFC. We used Dbh-Cre mice to express the inhibitory opsin Jaws^38^ specifically in noradrenergic neurons of the locus coeruleus and implanted an optic fiber above the injection site (**Fig. S1F**), and a second optic fiber was used to measure GRAB-DA. Mice were water restricted and GRAB-DA response to either a water reward or air puff in the mPFC while silencing noradrenergic neurons was measured (**Fig. S1G**). We found only a limited reduction in the GRAB-DA response to reward or air puff when the laser used for inactivating noradrenergic neurons was turned on (**Fig. S1H**). Therefore, our experimental approach allowed us to detect dopamine signals with suitable reliability and specificity.

### Broad dopamine signals to expected reward, but variable responses to omission

Head-fixed and water-restricted mice were trained over several sessions on a classical conditioning task in which a 1.5-second pure tone (the conditioned stimulus, CS+) was followed by a water reward (**Fig. 2A**). After several conditioning sessions, mice strongly associated the auditory cue with the reward, as characterized by rapid and sustained anticipatory licking during the cue (**Fig. 2B,C**). This anticipatory licking was significantly higher than the lick rate in response to a neutral cue (CS-), but comparable to consummatory licking following reward delivery (**Fig. 2B,C**). Additionally, anticipatory licking was similar across animals recorded in different brain locations (**Fig. 2D**).

**Figure 2.**
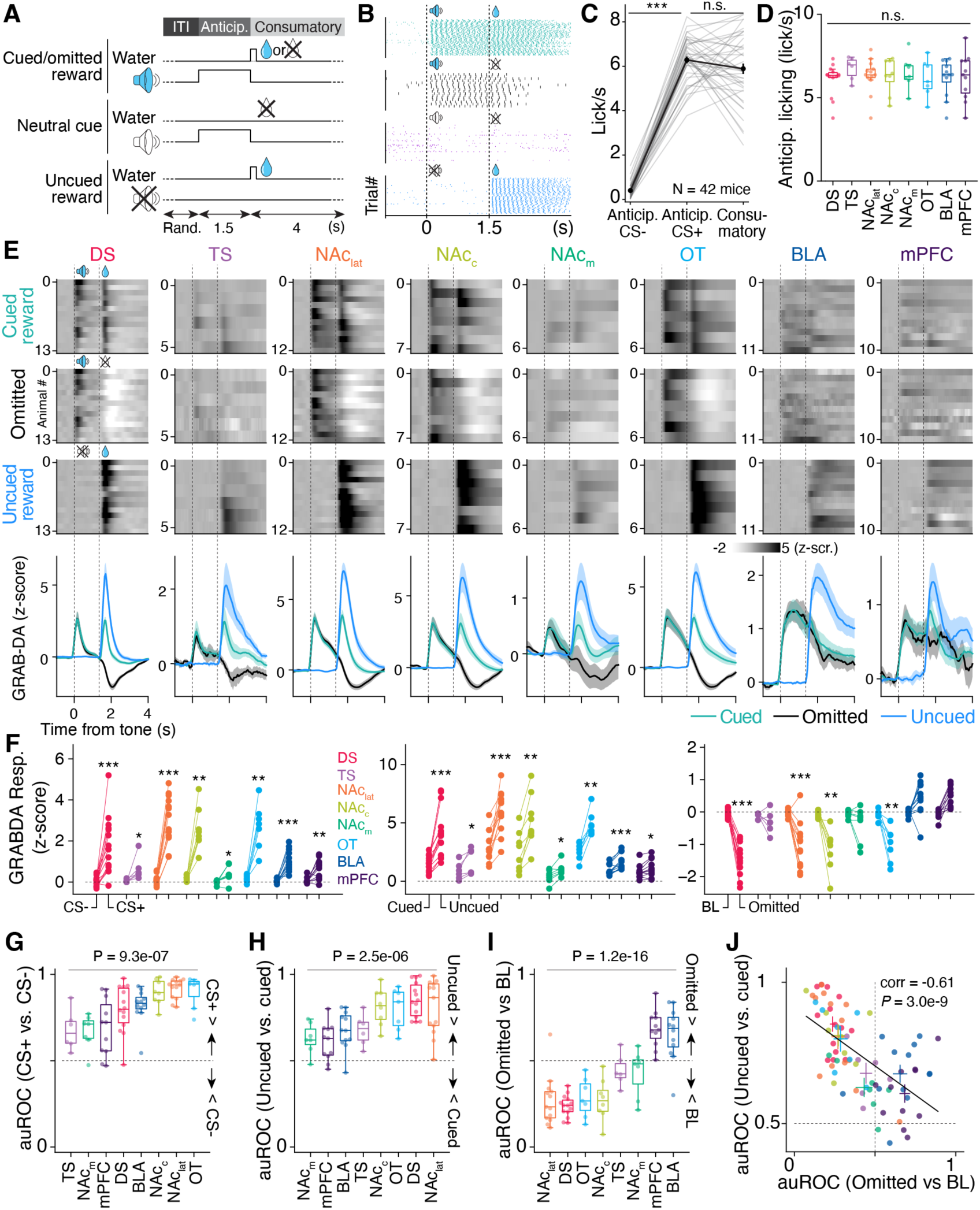
Broad dopamine responses to expected reward, but variable responses to omission. **(A)** Timing of inter-trial interval, tone, and reinforcement for four trial types. **(B)** Example licking behavior across all four trial types. **(C)** Average lick rate following the neutral (CS−) or conditioned (CS+) cue, or following water delivery. N = 42 mice. ***p < 0.001, Wilcoxon signed-rank test. **(D)** Distribution of anticipatory licking across different recording conditions. **(E)** Top: Raster plots of GRAB-DA signals from eight brain regions, averaged across animals, during cued, uncued, and omitted reward trials. Bottom: Population averages for each trial type in each region. GRAB-DA traces aligned to cue onset. Data shown as mean ± s.e.m. **(F)** Quantification of GRAB-DA responses in distinct brain regions for CS- vs. CS+ cues (left), cued vs. uncued rewards (middle), and baseline vs. omitted reward (right). Responses were calculated as area under the curve in a 1 s window post-cue or reinforcement. *p < 0.05; **p < 0.01; ***p < 0.001, Wilcoxon signed-rank test. **(G–I)** Area under the receiver operating characteristic (auROC) for CS− vs. CS+ (G), cued vs. uncued reward (H), and baseline vs. omitted reward (I). p values with one-way ANOVA. **(J)** Scatter plot showing the relationship between uncued vs. cued reward responses and omitted reward responses. Pearson r and p values shown. N = 13, 14, 8, 7, 6, 7, 11, and 12 recording sites for NAc_lat_, DS, NAc_c_, OT, TS, NAc_m_, mPFC, and BLA, respectively, in (E–J). Box-and-whisker plots in this figure, and for all figures in this article, plots show median, 25th–75th percentile, and min–max values.

Once mice learned the cue–reward association, we modified the task structure to include trials in which the reward was omitted following the cue, as well as uncued reward trials in which the reward was delivered without any cue (**Fig. 2A**). Using this paradigm, we recorded how dopamine release encoded reward prediction in various brain regions by comparing trials where rewards were either cued, uncued, or omitted (**Fig. 2E, Fig. S2A**). In all regions, we observed a significant dopamine response to the auditory cue associated with the reward (**Fig. 2E–G**). We also detected that dopamine release signaled the difference between expected and unexpected reward in all regions (**Fig. 2E,F**), with minor variability in encoding strength across targets (**Fig. 2H**). When comparing the kinetics of dopamine signals, we found that dopamine levels stayed elevated longer after reward delivery in some areas (**Fig. S2B,C**). For example, the dopamine signal in the mPFC or BLA persisted more than twice as long as in the DS (**Fig. S2D**). This was consistent with previous reports showing variable expression of the dopamine transporter (DAT) in distinct parts of the rodent dopaminergic system^39^. Therefore, our results show that dopamine encoding of reward is a conserved feature of the dopaminergic system, supported by distinct temporal dynamics.

When comparing dopamine signals to licking behavior, we noticed that in some brain regions – especially the mPFC and BLA – the temporal dynamics of dopamine release correlated strongly with the increase in lick rate following the conditioned stimulus (green traces in **Fig. 2B** vs**. 2E**). To examine the possibility that dopamine signals were linked to motor behavior rather than reward expectation, we quantified the correlation between lick rate and dopamine during anticipatory licking (**Fig. S2D,E**). We found limited differences in the correlation between anticipatory lick rate and GRAB-DA amplitude across regions (**Fig. S2E**). Furthermore, this variability in correlation only partially explained the variance in how dopamine encoded the cue (**Fig. 2G, S2F**). Hence, our recordings of dopamine signals primarily reflect reward expectation rather than motor output, consistent with recent findings indicating the dependence of learning on phasic dopaminergic release rather than movement initiation^40,41^.

In contrast to the relatively homogeneous dopamine responses to reward-predicting cues and unexpected rewards, we observed marked differences in the ability of dopamine release to signal reward omission across the dopaminergic system (**Fig. 2E,F,I**). Dopamine signals in the core and lateral shell of the NAc, the DS, and the OT showed strong suppression following reward omission after the conditioned stimulus, while other regions exhibited weaker suppression, or even signals above baseline in the BLA and the mPFC (**Fig. 2E,F,I**). This gradient in omitted reward encoding was linked to the dopamine response to unexpected rewards, where stronger unexpected reward signals correlated with a more negative dopamine response to omitted rewards (**Fig. 2J**). It was also linked with dopamine response kinetics, where slower dopamine release dynamics correlate negatively with omitted reward encoding signals (**Fig. S2G**). Together, our results suggest that dopamine release in response to omitted rewards differs by region, whereas the scaling of dopamine response to rewards by expectation is relatively consistent throughout the dopaminergic system.

### Dopamine responses to positive and negative reward expectations show regional variability

We adapted a variable reward task^42,43^ to examine how dopamine reward responses vary across different regions as a function of reward size and how expectation modulates this dopamine response. After training and testing on the classical conditioning task (**Fig. 2**), mice were exposed to cued or uncued reward trials in which the size of the reward was pseudo-randomly varied (**Fig. 3A**). Monitoring GRAB-DA during this task revealed that dopamine release increased monotonically with reward size (**Fig. 3B**). Across all recorded targets, dopamine responses significantly increased when comparing delivery of small (0.3 µL) to large (10 µL) rewards (**Fig. 3C**), although the strength of reward size encoding varied by region (**Fig. 3D**).We applied a Hill function to predict non-linearity of dopamine release as a function of reward size^43^ (**Fig. 3E, S3A,B**). This analysis revealed variability in the curvature of the reward response function across regions. In particular, the mPFC and BLA exhibited lower reward values required to elicit half-maximal dopamine responses (**Fig. 3F**). These results suggest that while dopamine encodes reward magnitude in all pathways, the curvature of the reward response function differs across targets of the dopaminergic system.

**Figure 3.**
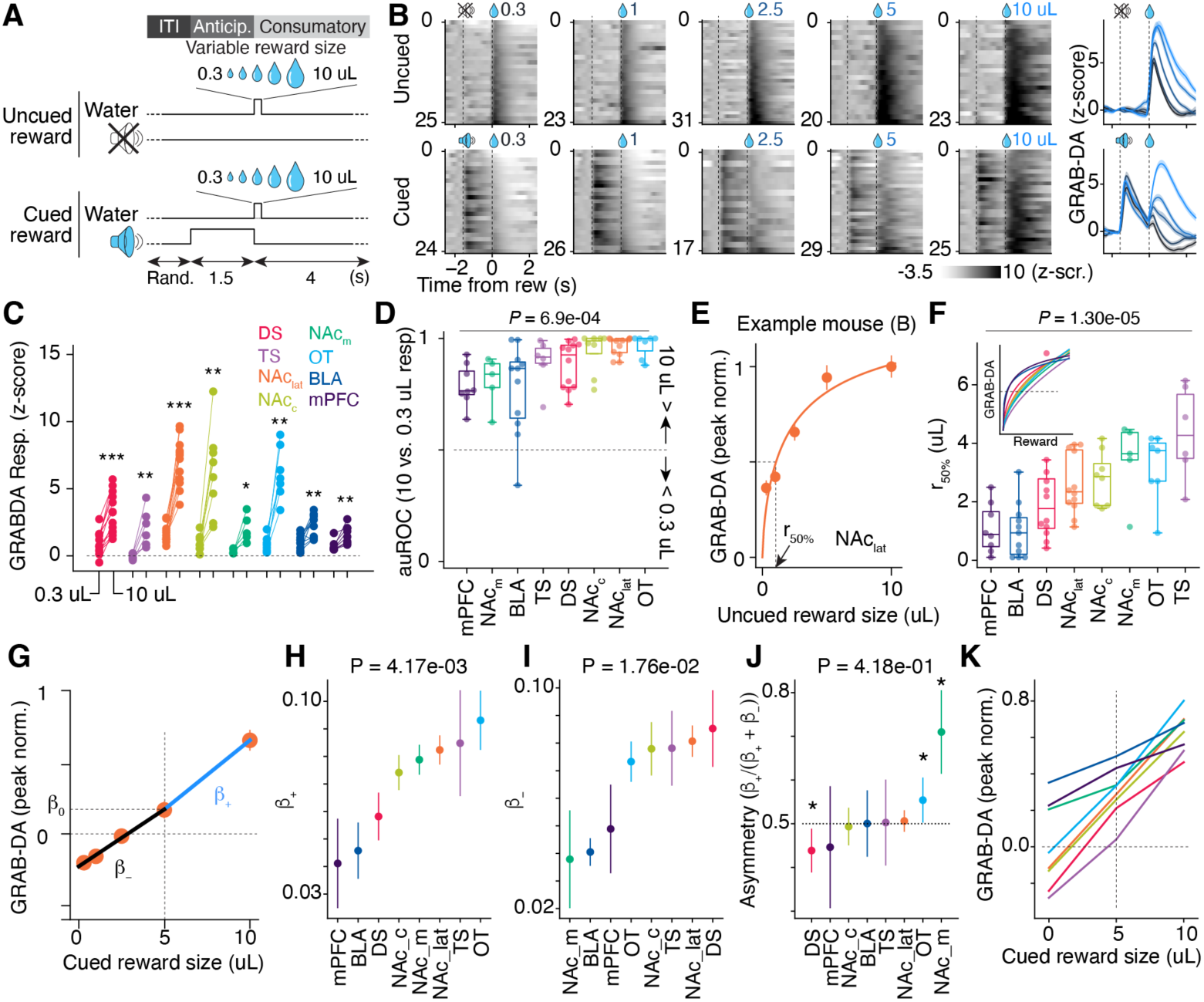
Dopamine responses to positive and negative reward expectations show regional variability. **(A)** Timing of cue and reward for the variable reward task. **(B)** Left: Raster plots showing GRAB-DA responses in an example session recorded in the NAc_lat_, aligned to the timing of water delivery for different reward volumes. Right: Session averages for different reward sizes for uncued (top) and cued (bottom) trials. **(C)** Average GRAB-DA responses calculated using a 1 s window following uncued 0.3 or 10 μL water reward. *p < 0.05; **p < 0.01; ***p < 0.001, Wilcoxon signed-rank test. **(D)** auROC comparing GRAB-DA responses to 0.3 and 10 μL uncued reward. **(E)** Average GRAB-DA responses to different volumes of uncued reward for the example shown in (B), peak-normalized. The solid line represents a least-squares linear fit of a Hill function (see Methods). The reward size eliciting a half-maximal GRAB-DA response is denoted *r_50%_*. **(F**) *r_50%_* values across eight recording sites. Inset: Mean GRAB-DA response as a function of reward size. **(G)** Mean GRAB-DA responses to cued reward volumes were fit with a piecewise linear regression constrained to intersect at the 5 μL reward point. Two linear regressions are shown for a representative NAc_lat_ recording. **(H–I)** *β+* represents the slope for rewards ≥5 μL; *β–* represents the slope for rewards ≤5 μL. **(J)** Asymmetry index, where values >0.5 indicate a steeper slope for larger-than-expected rewards. *p < 0.05, Wilcoxon signed-rank test. **(K)** Mean of all fitted piecewise linear regressions across regions using color codes from (H–J). Data are mean ± s.e.m. in (E, H–J). p values calculated via ANOVA in (D, F, H–J). N = 13, 6, 12, 8, 5, 7, 10, and 8 recording sites for DS, TS, NAc_lat_, NAc_c_, NAc_m_, OT, BLA, and mPFC, respectively.

Previous recordings of identified dopaminergic neurons have shown that the reward response function scales differently with expectation among individual dopamine neurons^42,44^, and distinct dopaminergic projections within the striatum^45^. To assess how reward expectation shifts the dopamine response as a function of reward size in distinct brain areas, we measured the average GRAB-DA signal following cued delivery of rewards of different sizes (**Fig. 3B,G**). To compare how dopamine signals scale with positive versus negative prediction error, we applied a piecewise linear fit centered on the mean GRAB-DA response to the cued 5 µL reward (**Fig. 3G**), the size used during conditioning. This piecewise linear model provided a better fit for cued responses than a single linear fit or a scaled Hill function (**Fig. S3C**), and allowed us to extract separate slopes for better- and worse-than-expected outcomes (**Fig. 3H,I; Fig. S3D**). Using this approach, we observed regional diversity in both positive and negative prediction error responses (**Fig. 3H,I**), along with some variation in asymmetry between these slopes (**Fig. 3J,K**). Dopamine release in the DS exhibited an asymmetric, steeper slope for negative prediction error, whereas in the NAcm and OT, the slope was steeper for positive prediction error (**Fig. 3K,L**). These findings suggest that reward prediction error signals may be differentially encoded across dopaminergic targets, potentially supporting region-specific learning dynamics.

### Region-specific encoding of an aversive stimulus by dopamine

Dopaminergic neurons can encode aversive events^5–7^, suggesting that variability in region-specific dopamine release supporting reward prediction error may reflect the signals link to aversive input. For example, if dopamine is released in a region following punishment, recordings from the same region could also show a positive response following reward omission. To test this, after recording from animals in the classical conditioning and variable reward tasks, we delivered uncued facial air puffs or reward. Air puffs are mildly aversive to mice and have been reliably used to suppress behavioral responses^46,47^.

As in the classical conditioning task, uncued reward delivery elicited increases in dopamine release across all recorded regions (**Fig. 2E**, **Fig. 4A,B**). However, dopamine responses to air puffs were absent in the OT, NAc_c_, and NAc_m_ (**Fig. 4A–C**). We did not observe valence-specific signaling – i.e., dopamine signals varying in opposite directions for reward versus aversive events – in any recorded region. When comparing responses to uncued reward versus air puffs, we observed larger dopamine increases to reward in the DS, NAc, and OT, whereas air puffs elicited stronger responses in the mPFC (**Fig. 4B,D**). We did not find negative correlation between dopamine responses to water reward and to air puffs (**Fig. 4E**), indicating that dopamine does not encode reward or punishment on the same axis in any of our recorded areas. In addition, we did not find a positive correlation between responses to air puffs (**Fig. 2I**) and to reward omission (**Fig. 4F**), nor to any other behavioral epochs linked with reward prediction error (**Fig. S4A**). This lack of correlation suggests that in regions like the DS or NAc_lat_, strong reward prediction error signals can coexist with robust responses to aversive stimuli. Altogether, these findings suggest high variability in how aversive events are encoded across the dopaminergic system. Moreover, our data indicate that canonical reward prediction error signals, in which dopamine would only signal reward or reward-predicting cues, may exist only in a subset of dopamine-targeted regions.

**Figure 4.**
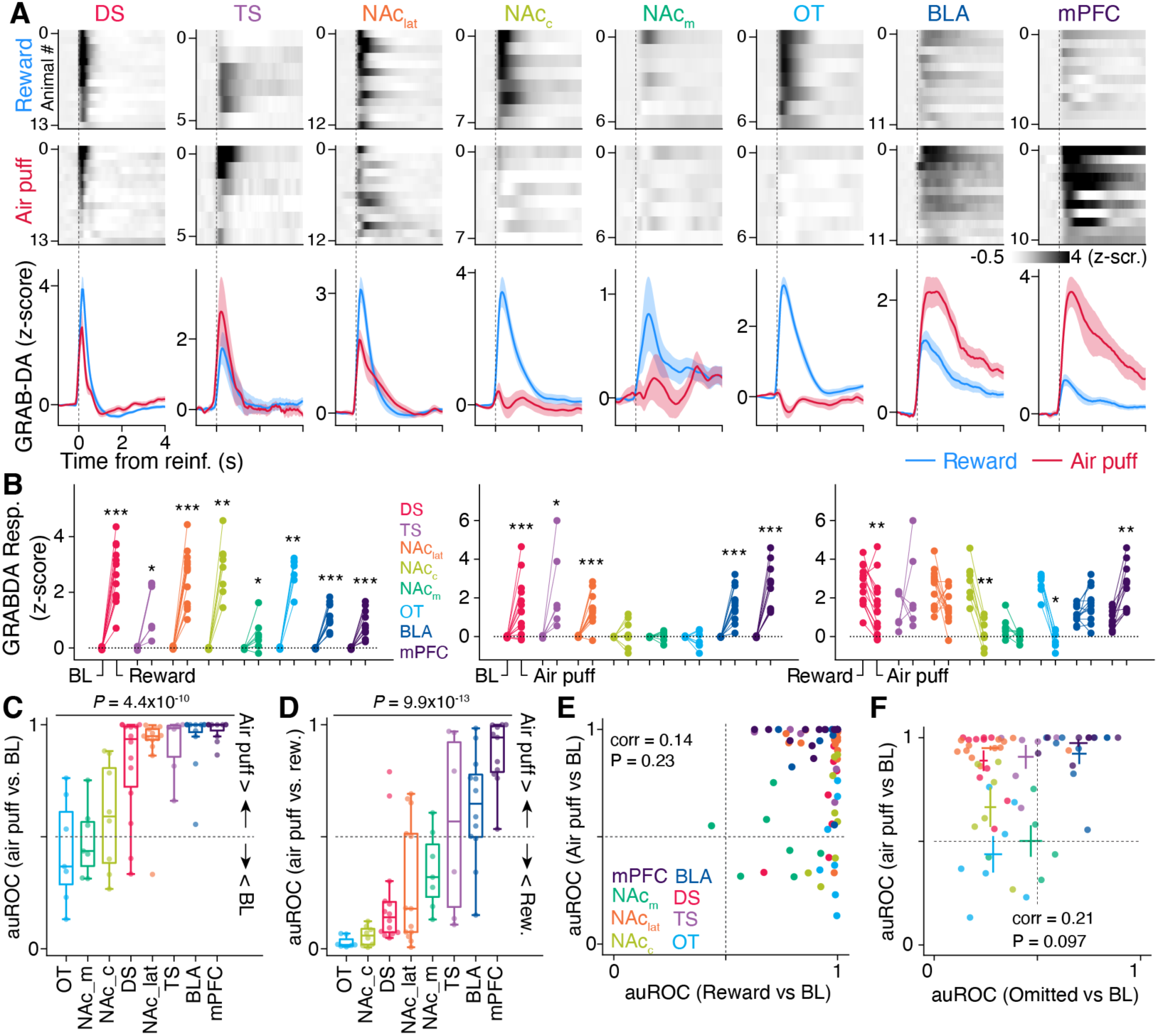
Region-specific encoding of an aversive stimulus by dopamine. **(A)** Top: Raster plots of GRAB-DA signals across eight brain regions, averaged by animal, for uncued reward and air puff trials. Bottom: Population average for the two trial types in each region. GRAB-DA signals were aligned to the timing of reinforcement. Data are plotted as mean ± s.e.m. **(B)** GRAB-DA responses in distinct brain regions to baseline (BL) and conditioned (CS+) cue (left), cued and uncued reward (middle), and baseline and omitted reward (right). Responses were calculated as the area under the curve over a 1 s window following cue or reinforcement onset. *p < 0.05; **p < 0.01; ***p < 0.001, Wilcoxon signed-rank test. **(C,D)** auROC comparing responses to air puff vs. baseline (C), and air puff vs. un-cued reward (D). p values calculated via ANOVA. **(E,F)** Correlation between air puff response and reward response (E), or omitted reward response (F). Correlation coefficients and p values were obtained using Pearson correlation. N = 7, 8, 14, 13, 7, 6, 12, and 11 recording sites from OT, NAc_c_, DS, NAc_lat_, NAc_m_, TS, BLA, and mPFC, respectively.

To assess whether variability in dopamine release to reward prediction error and aversive stimuli was dependent on recording location within each region, we calculated the correlation between individual dopamine response features and the anatomical location in a reference atlas of the fiber tip used for fiber photometry (**Fig. 1C, Fig. S4B-C**). We found limited correlation between fiber location and dopamine response magnitude within individual regions (**Fig. S4C**). However, when grouping data from the caudate putamen (CPu) and ventral striatum (VS), stronger correlations emerged between dopamine responses and recording site, suggesting that dopamine signaling may be organized along gradients within these larger anatomical divisions (**Fig. S4B-C**).

### Dopamine response patterns partially map onto recording location

Our findings thus far indicate a high degree of heterogeneity in dopamine signaling across brain regions during classical conditioning, variable reward, and aversive/rewarding events. We next asked whether these distinct patterns of dopamine responses to reinforcement are specific to individual targets of the dopaminergic system. If this were the case, we reasoned that dopamine response features could be classified according to their recording locations.

To test this, we analyzed data from mice with recordings across all three behavioral paradigms (**Figs. 2–4, S2–S3**) to determine if specific dopamine signals emerge from recording in specific dopaminergic pathways or brain regions (**Fig. 5A–C**). We applied Uniform Manifold Approximation and Projection (UMAP), as a dimensionality reduction technique, to visualize whether GRAB-DA response features clustered by recording site (**Fig. 5B,C**). By computing clustering quality, we observed poor separation between recordings from the caudate putamen (CPu) and ventral striatum (VS), whereas recordings from outside these major dopaminergic pathways – specifically the BLA and mPFC – clustered more distinctly (**Fig. 5B**). However, clustering was limited when analyzing individual recording locations (**Fig. 5C**), suggesting that dopamine response features only partially align with individual anatomical targets.

**Figure 5.**
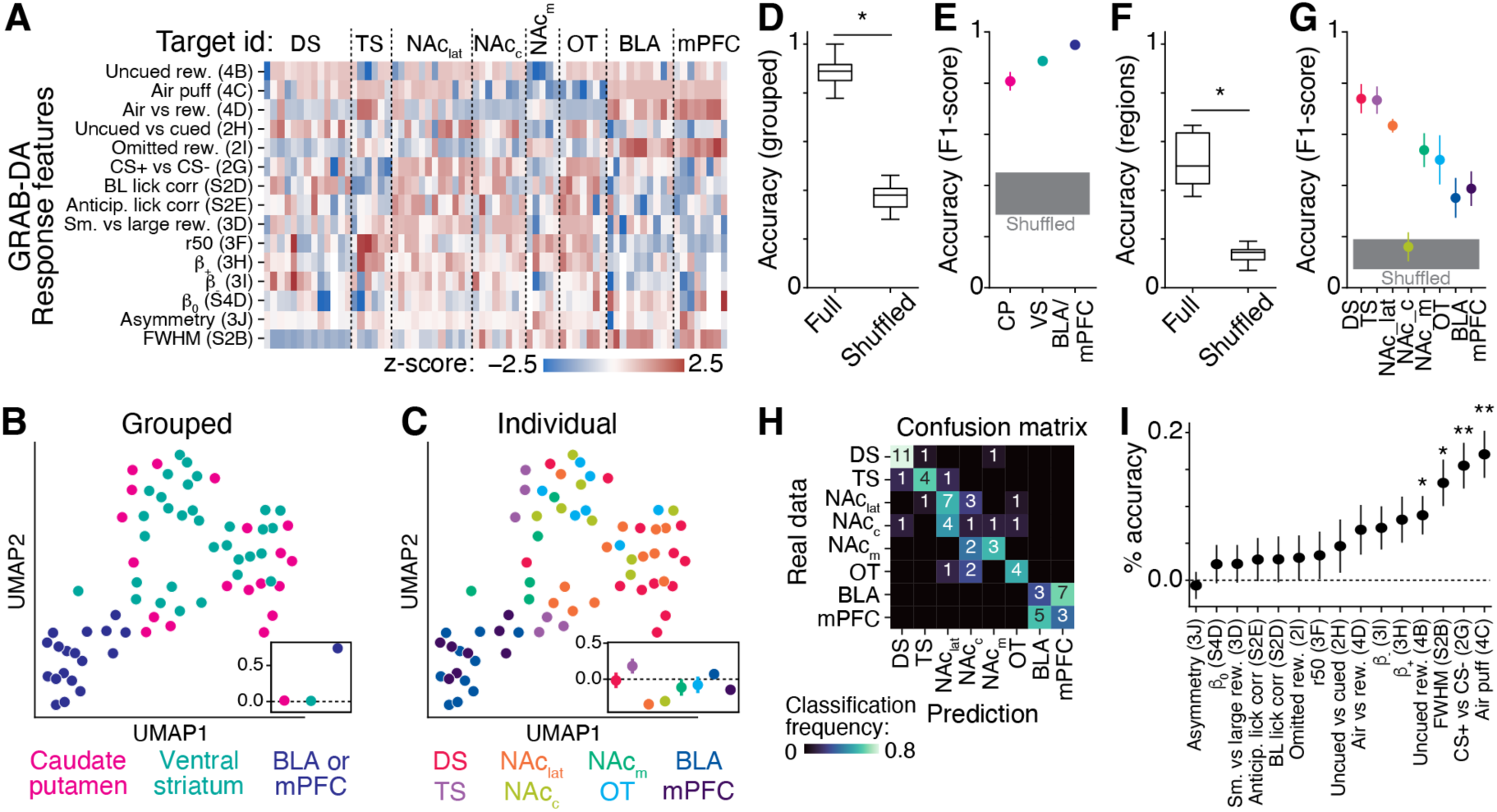
Dopamine response patterns partially map onto recording location. **(A)** Matrix of GRAB-DA response features measured across different conditioning tasks. Each column represents data from one animal, grouped by recording targets. Numbers in parentheses refer to corresponding figure panels. **(B,C)** UMAP projection illustrating the similarity of GRAB-DA response features shown in (A), color-coded by grouped dopaminergic pathways (B) or individual recording targets (C). Insets: Silhouette scores quantifying cluster separation for each label. **(D–I)** Support vector machine (SVM) classifier trained to identify the main dopaminergic pathways (D,E) or individual recording targets (F–I) based on GRAB-DA response features. Distributions of cross-validated classification accuracy are shown for both real and label-shuffled data. *p < 0.05, permutation test (D,F). (H) Cross-validated confusion matrix. (I) Relative contribution of each GRAB-DA response feature to classifier performance. *p < 0.05; **p < 0.01 Wilcoxon signed-rank test. Data are plotted as mean ± s.e.m. in (E), (G), and (I). N = 69 recording sites.

To further assess the spatial specificity of dopamine responses, we trained a support vector machine (SVM) classifier to predict recording location based on average GRAB-DA responses during classical conditioning, variable reward, and aversive/rewarding events. The classifier performed well when grouping data into the three major dopaminergic pathways (**Fig. 5D,E**; accuracy: 0.88 ± 0.03, *p* < 0.05 compared to shuffled data). Classification performance remained significantly above chance when decoding the eight individual target regions (**Fig. 5F,G**; accuracy: 0.52 ± 0.04, *p* < 0.05). Confusion matrix analysis revealed partial overlap in predictions between the ventral striatum subregions and BLA/mPFC recordings (**Fig. 5H**), consistent with their observed signal similarities. To identify which features most contributed to classifier performance, we performed permutation testing. The most important features were responses to air puff and CS+, as well as the full-width at half-maximum (FWHM) of the dopamine signal (**Fig. 5I**). Decoding accuracy remained above chance when using dopamine features from individual behavioral paradigms, although it was lower than when using the full feature set (**Fig. S5A–D**). Importantly, the variability in dopamine features across brain areas did not result from recordings in different mice, as simultaneous recordings within the same mouse showed similar variability (**Fig. S5E-G**). Together, our results demonstrate that dopamine response features to reinforcement events exhibit partial overlap across brain regions. Hence, while some anatomical specificity exists, the orchestration of dopamine signaling during reinforcement learning cannot be fully explained by anatomical segregation into distinct dopaminergic pathways.

### Distinct temporal trajectories of dopamine encode aversive and reward prediction errors

Thus far, our results reveal not only heterogeneity in dopamine signaling, but also lack of specificity in dopamine responses within the same brain area. For example, dopamine signals recorded in the NAc_lat_ resemble a reward prediction error both in classical conditioning (**Fig. 2**) and the variable reward task (**Fig. 3**). However, recordings from the same anatomical location in the same group of mice also showed responses to an aversive event (**Fig. 4**), whereas other regions, such as the OT or NAc_c_, showed minimal response to the same aversive stimulus. This discrepancy is further highlighted by the lack of correlation between dopamine responses to aversive and rewarding events (**Fig. 4E**), and between responses to aversive events and reward prediction error signals (**Fig. S4A**). To reconcile these findings, we considered the possibility that different dopaminergic pathways convey distinct components of the error signal^3,48^. This would mean that combining spatiotemporal dynamics of dopamine across brain areas may enhance differentiation between reward prediction error and other types of error linked with negative motivational signals.

We began by investigating how the temporal profiles of average dopamine signals correlate within and between different regions (**Fig. 6**). Specifically, we calculated Pearson correlation coefficients between average GRAB-DA signals recorded during key reinforcement epochs, including uncued and cued reward, air puff, reward omission, and reward-predicting cue (**Fig. 6A,B**). We found that dopamine signals were most strongly correlated across regions during unexpected reward and cue presentations, and least correlated during reward omission and the aversive stimulus (**Fig. 6C,D**). To further probe the circuit-level mechanisms underlying these correlations, we analyzed noise correlations – trial-by-trial co-fluctuations in dopamine signal strength – during simultaneous recordings in the mPFC, NAc_lat_, DS, and BLA (**Fig. 6E,F**). We focused on one-second epochs preceding stimulus onset (baseline) and following unexpected reward or air puff delivery. During baseline periods, significant positive correlations were observed for NAc_lat_-DS, mPFC-BLA, and NAc_lat_-BLA recording pairs (**Fig. 6G**). Unexpected reward delivery increased the number of significantly correlated region pairs, whereas air puff reduced them (**Fig. 6G**). These results suggest that trial-to-trial dopamine fluctuations are shared across a subset of dopaminergic targets, potentially reflecting shared dopaminergic inputs or coordinated activity of overlapping dopaminergic populations.

**Figure 6.**
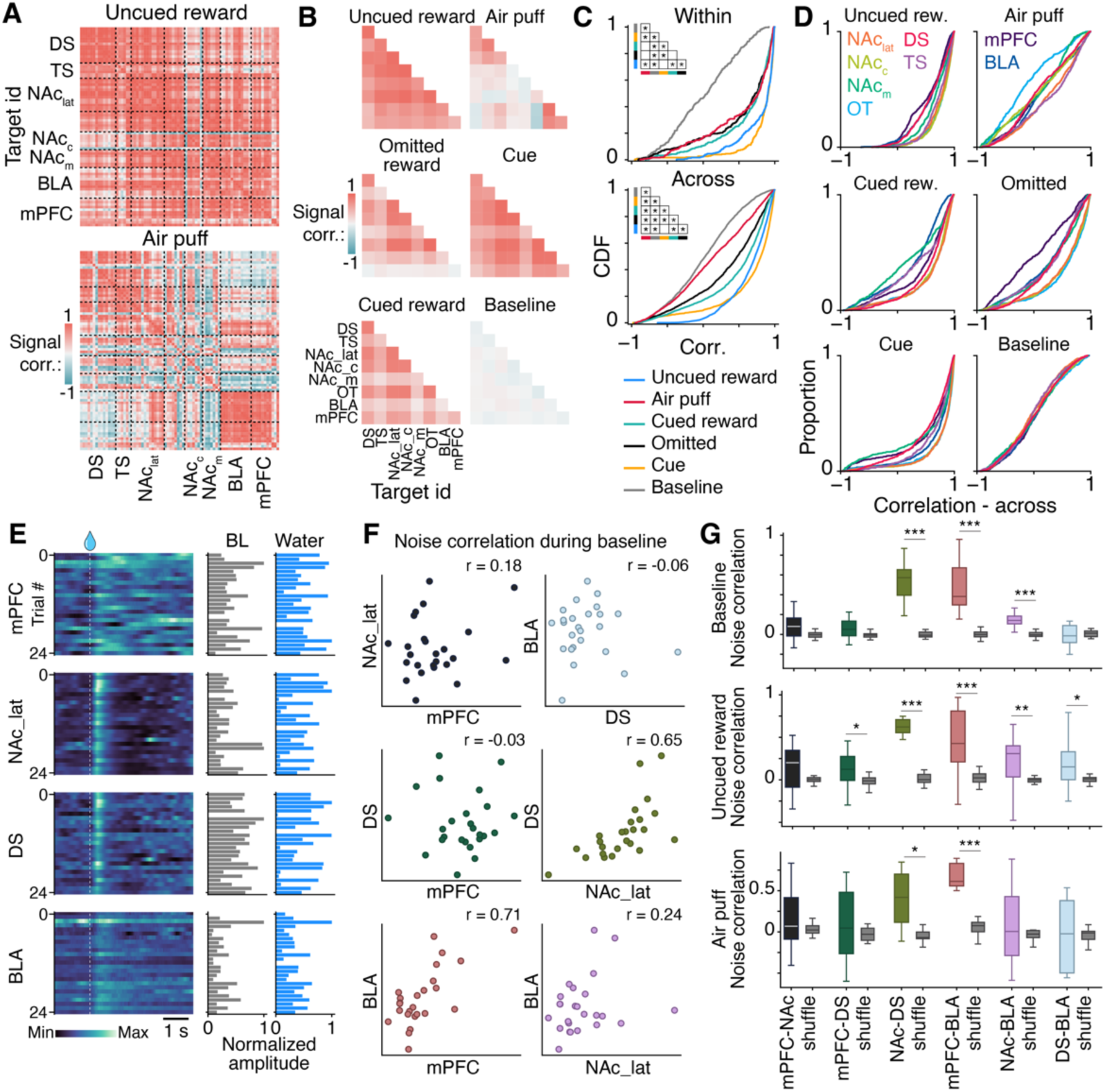
: Signal and noise correlation of GRAB-DA response to different behavioral events. **(A)** Correlation matrices of session-averaged GRAB-DA signals recorded in distinct brain targets. Matrices show pairwise correlations across individual mice during uncued reward (top) and air puff (bottom) conditions. Dashed lines separate data from different recording locations. N = 78 recording sites from 42 mice. **(B)** Mean pairwise correlation of GRAB-DA signals recorded in different brain locations. **(C)** Cumulative distribution function (CDF) plots of pairwise correlations across six different trial types. Top: within-target correlations; bottom: across-target correlations. Inset: P-values from ANOVA with post hoc Tukey HSD test; *: P < 0.05. N = 305 within-target and 2,628 across-target recording pairs. **(D)** CDF plots of across-target correlations for each behavioral epoch, separated by recording target. **(E)** Raster plots of GRAB-DA signals simultaneously recorded in mPFC, NAc_lat_, DS, and BLA in response to uncued reward. Right: average response during baseline and post-reward for each trial. **(F)** Correlation of baseline activity across simultaneously recorded GRAB-DA signals from the data shown in (E). Each dot represents one trial. **(G)** Noise correlations of GRAB-DA signals for baseline (top), uncued reward (middle), and air puff (bottom) epochs. N = 78 recording sites in (A–D); N = 28 sessions from 7 mice in (E–G). *: P < 0.05; **: P < 0.01; ***: P < 0.001 using a Wilcoxon signed-rank test compared to shuffled data with Bonferroni correction.

To test the hypothesis that different dopaminergic pathways convey distinct components of the error signal, we assessed how dopamine responses covary across our eight brain targets. We aligned mean temporal traces of dopamine release from these regions in response to key behavioral events and applied principal component analysis (PCA) to extract shared spatiotemporal patterns (**Fig. 7A**). We focused on the first three principal components, which together explained more than 97% of the variance (**Fig. S6A-C**). Visualizing these components in 3D space revealed clear separation of the air puff response from other reinforcement signals (**Fig. 7B**). This separation was particularly evident along the second principal component, which showed the greatest amplitude in response to the air puff compared to the reward-predicting cue and various reward conditions (**Fig. 7C**). To quantify this separation, we extracted the best-fit linear plane for each reinforcement trajectory in PCA space, effectively isolating the manifolds that constrain dopamine dynamics across events (**Fig. 7D**). By calculating the angles between the normal vectors of these planes, we found that the manifold associated with aversive events was significantly distinct from those associated with other reinforcement types. Notably, the plane corresponding to reward omission was not significantly orthogonal to those of reward-predicting cues or cued/uncued rewards. We further confirmed this spatial organization by analyzing responses to rewards of different magnitudes (**Fig. 7E**), showing that reward size scaled along PC1, the axis previously identified as representing reward prediction error. To evaluate the extent by which any combinations of multiple region-specific dopaminergic signals produces a low-dimensional separation of aversive from reward prediction error events, we tested how manifolds emerge in all possible combinations of three to seven recording locations (**Fig. 7F-H, S6D,E**). We found that separation of aversive from reward prediction error events increases with number of recording locations (**Fig. 7F, S6D,E**). Moreover, we found that combinations producing the largest low-dimensional separation included signals from DS, OT, mPFC and parts of the NAc (**Fig. 7G,H**).

**Figure 7.**
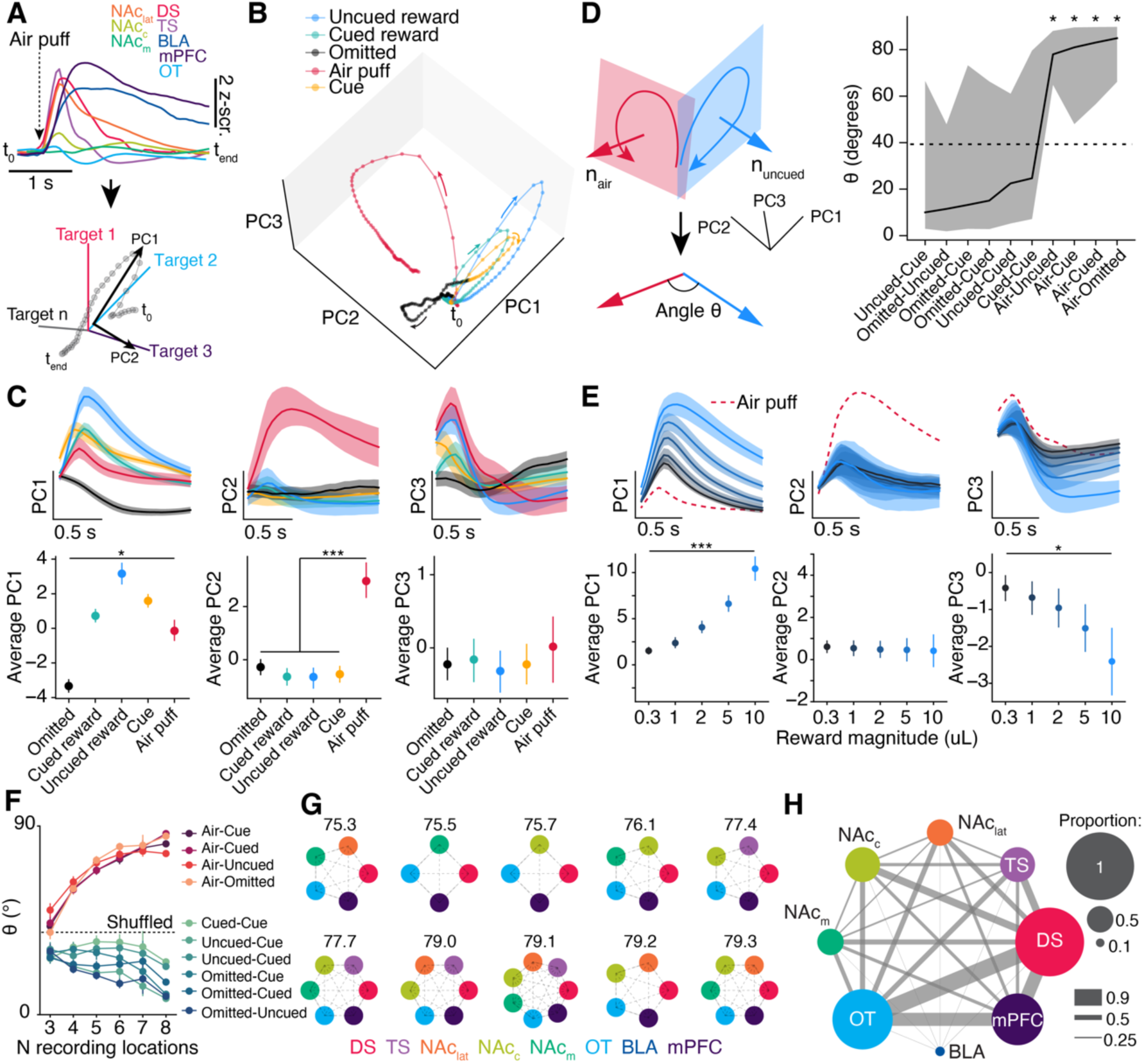
Distinct temporal trajectories of dopamine encode aversive and reward prediction errors. **(A)** Population average dopamine response to air puff (top), and schematic illustration of low-dimensional representations of dopamine dynamics (bottom). **(B)** Average trajectory in the first three principal components (PCs) during different behavioral epochs (B) **(C)** Top panels show the projection onto each PC; bottom panels show the corresponding average projections. Data are plotted as mean ± 95% confidence interval, calculated by bootstrapping the population average for each recording location. **(D)** A linear plane was fitted to each trial type, and the angle *θ* between the corresponding normal vectors *n* was computed for each pairwise comparison. The dashed line indicates the mean angle distribution for shuffled data. **(E)** Projection onto each principal component for uncued reward responses across different reward volumes; the air puff response is shown as a red dashed line for comparison. **(F)** Angle *θ* between manifolds calculated for each pairwise comparison as a function of the number of recording locations included for manifold computation. Data are mean ± sem, N = 56, 70, 56, 8, and 1 combinations of 3 to 8 recording locations. The dashed line indicates the mean angle distribution for shuffled data. **(G)** Top 10 combinations for which low-dimensional representations produce the largest differentiation of aversive from reward prediction error events. Difference between *θ_Air puff_* and *θ_RPE_* is shown above each combination. **(H)** Summary of the top 20 combinations producing the largest difference between *θ_Air puff_* and *θ_RPE_*. Dot size and line width represent the probability of a recording site or a recording pair respectively, being present in each combination. Data in (C-E) are plotted as mean ± 95% confidence interval, calculated via bootstrapping. N = 78 recording locations for (A-D, F-H) and 69 for (E). *p < 0.05; ***p < 0.001 in (C-E), Wilcoxon signed-rank test with Bonferroni correction.

These results indicate that while dopamine responses to aversive and reward prediction error events may co-occur in individual brain areas, their spatiotemporal covariation patterns form separable manifolds. This suggests that the dopaminergic system may distinguish reinforcement types through distributed population-level dynamics rather than localized signals alone.

## Discussion

Using multi-site recordings of dopamine release across the major anatomical divisions of the dopaminergic system, we provide a systematic characterization of how dopamine dynamics support reinforcement learning and how these signals are differentially distributed throughout the brain. Our results reveal that positive reward prediction error is a widespread feature of dopaminergic signaling, with regional differences in response strength, whereas aversive signaling appears more spatially segregated. Moreover, we show that this variability in dopamine response features, linked to both reward prediction error and punishment, partially align with specific projection targets. This supports the view that some dopamine signals are broadly distributed, while others are anatomically segregated. Finally, we demonstrate that brain-wide covariations in dopamine responses enable better dissociation of reward prediction error from punishment, suggesting that system-level dynamics across dopamine targets contribute to the computation of reinforcement signals.

During classical conditioning, we found that distinctive features of positive reward prediction error, such as cue responses and differences between expected and unexpected rewards, are broadcasted across the dopaminergic system. These observations align with previous studies of identified dopamine neurons in the VTA and SNc, which have shown widespread reward responses that are modulated by expectation^7,9,27,42^. Similarly, broad dopamine release in response to uncued reward and reward expectation has been reported in recordings across various basal ganglia subregions^18,20,22,24,28,49–51^. We also observed reward responses in the BLA and mPFC that scaled with expectation, consistent with dopamine recordings in those regions^29–31^ – though some studies report conflicting findings^32^. While previous reports suggest minimal reward-related dopamine activity in the tail of the striatum (TS)^23,25^, more anterior TS recordings have shown stronger reward responses^24^, in line with our observations. Additionally, reward magnitude modulates dopamine in the TS regardless of location^23,24^, which agrees with our results using variable reward sizes (**Fig. 3**). The cue and reward delivery have been shown to increase correlation among dopaminergic neurons^42,52^, and we similarly found increased across-site in dopamine trial-to-trial correlations in response to these events (**Fig. 6E-G**), suggesting that a shared input may underlie this system-wide broadcast of reward signals.

Conversely, our data show that dopamine does not uniformly encode worse-than-expected outcomes. Reward omission suppressed dopamine in the DS and parts of the ventral striatum (NAc_lat_, NAc_c_, OT), but not in extra-striatal regions like the mPFC and BLA, where levels remained above baseline. Most recordings of VTA dopamine neurons showed inhibition by reward omission^7,53^. Therefore, our findings suggest that non-inhibited dopamine neurons preferentially project outside the basal ganglia^29^. Additionally, slower dopamine reuptake in the BLA and mPFC (**Fig. S2B,C**), likely due to lower dopamine transporter expression^39,54^, could account for the lack of a detectable suppression in these areas (**Fig. S2G**).

Dopamine responses to aversive stimuli were highly variable across regions. Air puff stimulation failed to evoke responses in parts of the ventral striatum (OT, NAc_c_, NAc_m_), but elicited strong activation elsewhere. This variability mirrors what has been reported in single-unit^5–7,55^ and axonal^18,23,28,32^ recordings. Importantly, our dataset captures these dynamics across a broader range of dopaminergic targets than previously studied. We show that, although individual targets may display mixed responses to aversive and rewarding events, population-level covariation of dopamine signals across regions reveals distinct spatiotemporal signatures. This supports the idea that dopamine carries a vector-valued error signal rather than a scalar one, as previously proposed^3,48^. In this view, learning is driven not by a uniform signal but by distributed and potentially target-specific dopamine dynamics. Parallel processing of such information may be possible through the networks targeted by dopamine^56–58^, allowing for more selective forms of learning and behavior modulation.

Finally, although fiber photometry offers a powerful tool for estimating average dopamine release within a localized region (∼hundreds of microns), it may underrepresent finer spatial heterogeneity of dopamine signaling. Dopamine neurons can be molecularly subdivided based on gene expression profiles^12^, and while these subtypes often project to different targets, their projection fields can also overlap within the same region^11,49,59^. However, dopamine signals measured using fiber photometry likely reflect an average across these subtypes, potentially preventing the measurement of functional differences. Imaging of individual dopamine axons in the DS and mPFC has shown marked diversity in how different axons encode rewarding, aversive, or motor signals^29,49,60^. Moreover, imaging of dopamine release in the dorsal striatum using two-photon microscopy has revealed wave-like dynamics in response to reward^51^. Dopamine acts via both synaptic and volume transmission^61^, raising questions about how local fluctuations in dopamine levels relate to neuromodulatory signaling at the cellular level. Future work incorporating higher spatial resolution and genetic dissection of dopaminergic circuits, while maintaining a system-wide approach, will be essential to further resolve these complexities.

## Resource availability

### Lead contact

Request for further information or resources will be fulfilled by the lead contact, Vincent Breton-Provencher.

### Materials availability

This study did not generate new reagents.

### Data and code availability

Custom python scripts used for analyzing the data, controlling the behavior, and registering coronal slices to the Allen Brain mouse reference atlas will be made available at the time of publication.

## Acknowledgements

We thank Lea-Maude Gauthier and Catherine Bédard for behavioral training. Antoine Légaré for feedback on some of the analyses. Dr. Yulong Li (Peking University) for providing early access on the GRAB-DA3m sensor. Dr. Grayson Sipe and Dr. Paul Masset for valuable feedback on the original manuscript. Support to the VBP lab was provided by NSERC Discovery Grant (RGPIN- 2021-03284), CIHR project grant (#517536), CFI John R. Evans Leaders Fund (Grant No. 44014), AFOSR Cognitive & Computational Neuroscience Program (FA9550-23-1-0533), a Future Leaders in Canadian Brain Research Program from Brain Canada, and a Research Scholars— Junior 1 Salary Award from Fonds de recherche du Québec (FRQ), Santé. Support to ML lab was provided by NSERC Discovery Grant (RGPIN-2024-05363), CIHR project grant (#451548) and Research Scholars—Senior Salary Award from Fonds de recherche du Québec (FRQ), Santé and Parkinson Quebec.

## Author contributions

Conceptualization, formal analysis, writing – original draft: S.J.B. and V.B.P.; Methodology and investigation: S.J.B., J.B., V.B.P.; Resources, supervision, funding acquisition, writing – review & editing: M.L., V.B.P..

## Methods

### Animals

All animal procedures were approved by Université Laval’s Animal Protection Committee and conformed with the guidelines established by the Canadian Council on Animal Care. Two to six months old male and female mice were used in this study. Mice were housed in a room with light on from 7 am to 7 pm in ventilated cages with controlled humidity (50 to 70 %) and temperature (22 to 25 °C). Food and water were provided ad libitum unless noted (see *Behavioral training and task*). All experiments were performed during the light part of the cycle. Most experiments were carried out in C57Bl6 wild-type mice. The *Dat-IRES-Cre* line (B6.SJL-Slc6a3tm1.1(cre)Bkmn/J; JAX stock #006660) was used for specific expression of the activating opsin Chrmine2.0 in dopamine-expressing neurons of the substantia nigra or ventral tegmental area (**Fig. S1C-E**). The *Dbh-Cre KI* line (B6.Cg-Dbhtm3.2(cre)Pjen/Jl; JAX stock #033951) was used for specific expression of the silencing opsin Jaws in noradrenaline-expressing neurons of the locus coeruleus (**Fig. S1F-H**).

### Surgeries

Mice were anesthetized with isoflurane (4% induction, maintained with 1.5-2%). General analgesia (carprofen, 2 mg/ml) and local anesthesia (lidocaine/bupivacaine, 0,8 mg/ml – 0,4 mg/ml) were given before the beginning of the surgery. Ringer’s lactate solution was injected at the beginning of the surgery and at every hour. Following deep anesthesia, mice were placed in a stereotaxic frame, their head was shaved and cleaned, and their scalp was removed with surgical scissors. The skull was aligned in the stereotaxic frame (Model 942, Kopf) by leveling the heights of the bregma and lambda. A virus expressing GRAB-DA3m ^33^ (AAV2/9-hSyn-GRAB-DA3m, titer of 5.51 x 10^12^ GC/mL, BrainVTA) was injected in one or more of the following targets : DS [antero-posterior (AP) : 0.2-0.8 mm, medio-lateral (ML): 1.7-2 mm, dorso-ventral (DV): -2.5 mm], TS [AP : -1.8, ML : 3.25, DV : -2.5], NAc_lat_ [AP : 1.57, ML : 1.65, DV : -4.4], NAc_c_ [AP : 1.57, ML : 1.15, DV : -4.2], NAc_m_ [AP : 1.57, ML : 0.45, 19 DV : -4], OT [AP : 2, ML : 1, DV : -4.2], BLA [AP : -1.6, ML : 3.35, DV : -4.95], and the prelimbic division of the mPFC [AP : 2.1, ML : 0.3, DV : -1.2]. 200 nL of the virus was injected at a volume rate of 100 nL/min using a nanoliter injector (NANOLITER2020, WPI inc). Next, a fiber optic cannula (0.39 NA, 400 um, RWD) was implanted 0.2 mm above each injection site for fiber photometry measurements. A custom head plate, which was used for head fixation, was positioned on the skull of the mouse. The cannulas and the head plate were attached to the skull with adhesive cement (C&B Metabond, Parkell). For optogenetics experiments in which we activated dopamine-expressing neurons, we injected 400 nL of a cre-dependent virus coding for the opsin ChRmine 2.0 (AAV2/8-nEF-Con/Foff 2.0-ChRmine-oScarlet, titer of 2.1 x 10^13^ GC/mL, #137161-AAV8 Addgene) in the VTA [AP : -3.3, ML : 0.4, DV : -4.2] or the SNc [AP : -3.5, ML : 1.2, DV : -4] of *Dat-Cre* mice. For optogenetics experiments in which we silenced noradrenaline-expressing neurons, we injected 400 nL of a cre-dependent virus coding for the opsin Jaws (AAV2/5-CAG-Flex-rc Jaws-KGC-GFP- ER2, titer of 1.3 x 10^13^ GC/mL, #84445-AAV5 Addgene) in the LC [AP : -3.3, ML : 0.4, DV : -4.2] of *Dbh-Cre* mice. A fiber optic cannula (0.39 NA, 400 um, RWD) was also implanted 0.2 mm above the opsin injection site. Animals were kept under general analgesia for 48h following surgery. Mice were single housed after surgery for improved recovery and for handling water regulation (see *Behavioral training and task*).

### Behavioral apparatus

Mice were head-fixed on a behavioral rig and confined in a plastic tube (a modified 50-mL centrifuge tube) to restrict body movements. The behavioral rig was adapted from a previously published set up^46^. Briefly, a metallic lick spout, placed ∼2 mm away from the mouth and connected to a custom-made lick detector, was used to deliver water rewards. Rewards of various size (0.3 uL to 10 uL) were delivered from a 3 mL water reservoir positioned above the behavioral rig and connected to a solenoid valve (MB2029-036B-B-Y1-203, GEMs sensors). Durations of valve opening were calibrated and tested routinely to provide a precise and reliable delivery of reward volumes. A small tube at 3 cm from the mouse facial area was used to deliver air puff punishment (compressed air at 50 psi for 0.3 s). The tube was connected to a Picospritzer III (Parker Hannifin) for controlling the timing and repeatability of air puff pulses. A microcontroller board (Arduino Nano Every) was used to control the timing of reward/air puff delivery, to record voltage signals from the lick detector, and to send a TTL signal at the end of each trial for syncing with the fiber photometry system. Sound stimuli (1.5 s duration, 4 or 12 kHz) were delivered via two speakers positioned 25 cm from each side of the mouse. The sound stimulus intensities were established by a sound level meter. The Arduino microcontroller controlling and recording the behavioral setup was connected to a computer running a custom-written Python (version 3.11 or 3.12) script that was able to record lick rate, while controlling the timing of the sound cue (using the Simple Audio library), water, and reward. Behavior rigs were assembled primarily with optomechanical components (Thorlabs). Behavior was run inside a foam-lined sound attenuating cubicle (ENV-022, Med Associates).

### Behavioral training and task

At least a week after surgery, mice were water restricted by progressively reducing their daily water intake (from 2.5 mL to 0.8-1.5 mL). During water restriction, the weight of each mouse was maintained above 80% of its initial weight, and body condition score was monitored closely. Mice were habituated to head fixation and behavior rig for 5-10 min during 2 consecutive days before the start of training. During this habituation period, a few water rewards were delivered from the lick spout to initiate licking and to reduce stress.

Each trial started with an inter-trial interval randomly drawn from an exponential distribution (mean: 10 s, range: 0 to 45 s). This was followed by an auditory pure tone of 1.5 s duration. A reinforcement - a drop of water (5 uL for the classical conditioning task or 0.3-10 uL for the variable reward task), an air puff, or nothing - was delivered at the offset of the tone. The end of each trial was followed by a fixed delay period of 4 s. We used a pseudo-randomized trial sequence for all behavioral training and testing. We ensure with this pseudo-randomized order that the proportion of each trial type presented was similar for every block of ten trials.

During training, we used two types of trials: conditioning trials where a 12 kHz auditory cue (the conditioned stimulus, CS+) was immediately followed by a 5 uL water reward and trials where only a 4kHz tone was presented (neutral cue, CS-). Training sessions consisted of an average of 80 trials per day and lasted until the animal was fully conditioned on the 12kHz auditory tone.

Conditioning was evaluated by comparing the rate of anticipatory licking during the CS+ and consummatory licking following water delivery. A mouse was considered trained if these two rates were nearly equal (**Fig. 2C**). Training sessions lasted for a minimum of 10 sessions and until the mouse was fully conditioned. We used a tone intensity of 35 dB for each cue, calculated by measuring the sound pressure level for either CS+ or CS- cue and subtracting the noise level of that given frequency.

Following training, we recorded the dynamics of dopamine release in mice during the classical conditioning task. In this task, we omitted the reward following the CS+ on a fraction of trials. We also added trials in which a water reward was delivered without any predicting cue. The sequence of trial types was randomized using these probabilities: 0.4 of CS- trials, 0.36 of cued reward, 0.04 CS+ with omitted reward, and 0.2 of uncued reward. Mice typically performed an average of 108 ± 2 trials per session on the classical conditioning task. Each mouse was recorded for four sessions on the classical conditioning task before switching to the variable reward task. In the variable reward task, all trials were rewarded but with reward volume randomly chosen between 0.3, 1, 2.5, 5 or 10 uL. Trials were equally divided into either cued or uncued reward delivery. Mice typically performed an average of 165 ± 5 trials per session on the variable reward task and were recorded for two sessions. On the last recording sessions, we measured dopamine release dynamics in response to aversive or rewarding stimuli by randomly delivering air puffs to the facial area of the mouse on 20% of all trials. For these sessions, we also included un-cued reward on 20% of the trials. Note that cued reward trials were also included in these sessions but not analyzed.

### Multi-site fiber photometry

Fiber photometry recordings were done at least 4 weeks post-surgery. We used a custom-made fiber photometry system adapted from a previously published design ^32,62^. Briefly, the light from two mounted LEDs of 415 and 470 nm wavelengths (#M415L4 and #M470L5, Thorlabs) were focussed with aspherical lenses (#LA1951 – N-BK7, Thorlabs), bandpass filtered (#FB410-10 and #FB470-10, Thorlabs), and combined with a dichroic mirror (#DMLP425R, Thorlabs). The emission light was then reflected using a dichroic mirror (#FF495-Di03-25x36, Semrock) onto the back aperture of a 10X objective (UPlanXApo 0.40NA/3.1WD) used for collimating the light onto the single optical connector side of a multi-fiber optic patch cord. We either used two-ended (SBP(2)_200/220/900/900-0.37_1m_FCM-2xFC, Doric) or four-ended (BBP(4)_400/440/900- 0.37_2m_FCM-4xMF1.25_LAF, Doric) low-autofluorescence patch cords to simultaneously record from two or four targets. Emission light was collected back onto the microscope objective, bandpass filtered (#FF01-535/22-25, Semrock), and detected with a CMOS camera (Blackfly S BFS-U3-04S2M, Teledyne). To isolate the isosbestic from the GRAB-DA signal, the 415 and 470 were alternated at a rate of 40 Hz for a final acquisition rate of 20 Hz. We set the power at the output of the patch cord to 35-50 uW for the 470 nm light. Once the patch cable was connected to the optic fiber cannula on the animal, the power of the 415 nm LED was adjusted to detect an emission power equivalent to the 470 nm signal. Synchronization of the camera frame rate and the two LEDs was done by a microcontroller (Arduino Nano Every). A second microcontroller (Arduino Uno) was used to synchronize the fiber photometry system with behavior and to interface the system with a computer for data acquisition. A custom Bonsai script was used for saving raw fluorescence signals from individual optic fibers imaged onto the camera as well as the TTL signal from the behavior.

Raw fiber photometry data were processed with a custom-made Python script. The isosbestic and GRAB-DA signals were separated, and each signal was high-pass filtered at 0.005 Hz using a 2^nd^ order Butterworth filter to remove slow drifts in fluorescence intensity. To correct for motion or light artifacts, the isosbestic signal was subtracted from the GRAB-DA signal. The signal was then smoothed with a 5-point moving average. Using the onset of the TTL signal, the corrected GRAB-DA signal was then aligned to the beginning of each trial. The baseline period for each trial was defined by a 0.5-second-long window preceding the tone onset, or reinforcement when no tone was presented. GRAB-DA signal was z-scored by dividing by the standard deviation of the baseline signal. The GRAB-DA signal for each trial was baseline subtracted except for noise correlation calculations (**Fig. 6E-G**).

### Optogenetics

For activating dopamine expressing neurons and record from either the NAc_lat_, DS, or the mPFC, we used a LED light source of 590 nm (M590F3, Thorlabs) wavelength controlled by a microcontroller (Arduino UNO). We used a power of 3.15 mW, measured at the tip of the patch cord. The patch cord was connected to the optic fiber cannula positioned above the VTA (for NAc or mPFC experiments) or the substantia nigra (DS) of a head fixed mouse. Trains of 10 ms light pulses at a frequency of 10 Hz were presented every 15 seconds. We used 1 (0.1 s), 2 (0.2), 5 (0.5), 10 (1) or 20 (2) light pulses, and each pulse train were repeated 30 times. For silencing noradrenergic activity from the locus coeruleus and measuring GRAB-DA signals in the mPFC in response to water or air puff delivery, we used a laser diode light source of 638 nm wavelength ( LDFLS_473/070_638/120, Doric Lenses). We used a power of 15 mW, measured at the tip of the patch cord. The patch cord was connected to the optic fiber cannula positioned above the LC of a head fixed mice. Trial sequence alternated between reward, air puff, light on only, light on+reward, and light on+air puff. Each trial type was repeated 15 times using a 20 second inter-trial interval. We used a continuous light pulse that was started 1 second before reward or air puff delivery, and lasted for 3.5 seconds, including a 0.5 second ramp down of the light power. For analyzing the GRAB-DA response in experiments in which we photoactivated DA-expressing neurons, we baseline-subtracted each trial and calculated the area under the curve for a period of 1 s following a single 10-ms pulse of light in the VTA or SNc. For analyzing the GRAB-DA response in the mPFC when we photoinactivated noradrenergic neurons of the locus coeruleus, we baseline-subtracted each trial using a period preceding the air puff or water delivery. We then calculated the area under the curve for a period of 1 s following the air puff or water delivery and compared laser on versus off trials.

### Histology

Mice were anesthetized with ketamine/xylazine (10 mg/ml; 1 mg/ml), perfused with standard phosphate buffer solution (PBS, P4417, Sigma-Aldrich), and then perfused with a 4% para formaldehyde solution (PFA, #15710, Electron Microscopy Science). Following perfusion, brains were immersed in 4% PFA for 24 h for post-fixation and then kept at 4°C in PBS. Fixed brains were sliced coronally at a thickness of 100 μm using a vibratome (Leica VT1000S). For some of the slices, we confirmed the expression of GRAB-DA by staining for GFP with immunohistochemistry. These slices were blocked and permeated at room temperature in a 10% NGS, 0.5% triton solution for 1h, then incubated with the primary antibody anti-GFP (1:500, rabbit, Synaptic systems, 132 002) at 4°C for 12h-24h. The next day, the slices were washed with a 1% NGS, 0.05% triton solution and were incubated with the secondary antibody AlexaFluor 555 anti-rabbit (1:400, Invitrogen, A11011) for 2h at room temperature. Slices were mounted using Prolong Gold Antifade Mountant with DNA stain DAPI (P36931, ThermoFisher) and imaged on a Axio imager M2 epifluorescence microscope (Zeiss) or on a confocal microscope (LSM700, Zeiss). All images were processed with the open-source software ImageJ^63^ to adjust brightness and pixel resolution. Anatomical position of the fiber tip was registered to the Allen Mouse Brain Common Coordinate Framework (CCFv3)^34^ using a custom-made Python script. This script enabled us to browse the atlas in the coronal axis and title the coronal plane angle to find the best section of the atlas that matched our brain slice. Then the slice was registered onto this atlas section using point registration. Fiber registration confirmed placement in one of the eight selected targets, and allowed for calculating the correlation between fiber location and GRAB-DA response (**Fig. S4**).

### Data analysis

All data analysis, unless noted, was performed using custom-made scripts written in Python (version 3.9 to 3.12).

To quantify GRAB-DA responses, we calculated the area under the curve (AUC) following stimulus onset. A 0.5-second window was used for auditory cue, cued reward, omitted reward, uncued reward, and air puff responses. A 1-second window was used for variable reward response calculations. Decoding performance was assessed using receiver operating characteristic (ROC) curves, calculated by comparing GRAB-DA amplitude distributions across trials. The area under the ROC curve (auROC) was used to indicate decoding performance, with auROC values close to 0.5 representing chance-level performance. For analyses of conditioned auditory stimuli, we included parts of behavioral sessions where the animal exhibited a lick rate of at least 3 licks/s following the CS+ stimulus (averaged over 5 trials).

Dopamine responses to uncued rewards of varying sizes were fit using the Hill function^43^: *f*(*r*)=1/(1+(*σ*/*r*)^e^), where *r* is the reward size. The parameter *σ* was fit to the peak-normalized data using least-squares regression. The Hill coefficient *e* was determined empirically as the value yielding the maximum coefficient of determination (R²) (0.35 ± 0.03). The reward size producing half the GRAB-DA response (*r_50%_*) was calculated from the fitted Hill function *f*(*r*). Responses to cued rewards of varying sizes were fit using a piecewise linear function:

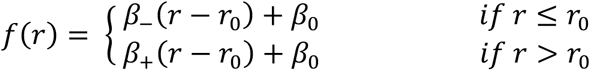

where *r* is the reward size, *r₀* is the reward size used during conditioning (5 µL), *β_-_* and *β_+_* are the slopes, and *β_0_* is the common intercept. The asymmetry of the piecewise linear function was determined for each recording location using *β_-_* / (*β_-_* + *β_+_*). We also tested a simple linear regression and a scaled Hill function, where the function output was subtracted by an expectation factor. The piecewise linear function provided the best fit, followed by the scaled Hill function.

Classification of recording locations based on GRAB-DA response features acquired during various behavioral experiments was performed using dimensionality reduction. For this analysis, we included recordings from brain regions sampled across all three behavioral paradigms (classical conditioning, variable reward, and reward/aversive) in the same animal. We employed the ’umap-learn’ (version 0.5.7) library to perform UMAP dimensionality reduction of our GRAB-DA feature data and evaluated the clustering quality of each recording location using the silhouette score. We also compared our results with PCA and t-SNE dimensionality reduction methods, obtaining similar outcomes. In a separate analysis, we trained a support vector machine (SVM) classifier to predict recording location based on GRAB-DA features across all behavioral paradigms using the Scikit-learn library for Python (version 1.3.0). We assessed the classification accuracy (% of correctly predicted locations) using 8-fold cross-validation, ensuring that data from each recording location was included in both training and testing folds. A permutation test (n=25) was used to evaluate the statistical significance of the accuracy. We assessed the classification performance for individual recording locations using the F1 score: 2\**TruePositive / (2\**TruePositive + FalsePositive + FalseNegative). The contribution of each feature to the SVM classifier was determined by measuring the decrease in accuracy after shuffling the values for a feature (repeated 100 times). Additionally, we tested other classification methods, including random forest, logistic regression, and naive Bayes, and observed slightly lower classification accuracies: 0.45, 0.43, and 0.48, respectively, compared to 0.52 for the SVM method.

To measure signal correlation, we calculated Pearson’s correlation between average GRAB-DA responses for each recording location, both within (e.g., DS_#2_ vs. DS_#1_) and across (e.g., DS_#1_ vs. TS_#1_) targets. Signal correlation was determined using a 2-second window (-1 to 1 s) for auditory cue-linked activity and a 5-second window (-1 to 4s) for reinforcement-linked activity. To measure noise correlation, we used data from simultaneously recorded animals in the DS, NAc_lat_, BLA, and mPFC. Trial-by-trial GRAB-DA response amplitudes were calculated using a 1-second baseline preceding or following air puff or water delivery. Pearson’s correlation was calculated for each pair of simultaneously recorded responses. Significance was evaluated by comparing the correlation to a null distribution generated from shuffled trial sequences (average of 10 shuffles per pair).

To analyze system-wide dopamine dynamics related to reward prediction error and aversive stimuli, we visualized time-dependent activation of all eight recorded signals using dimensionality reduction. A window of 0.1 to 1.3 seconds after the conditioned cue, neutral cue, omitted reward, cued reward, and uncued reward was extracted from the population average for each target. These signals were concatenated into a matrix of 8 targets by 168 time points. We used principal component analysis (PCA) to reduce dimensionality, retaining the first three components which explained 97% of the variance. These components were used to transform the data from variable reward experiments. Manifolds representing different behavioral epochs were estimated using the equation: *pc_3_* = *m*\**pc_1_ + n*pc_2_* + *c*, where *pcᵢ* are the principal components. Plane normals, defined with the vector: [m, n, -1], were defined using least-squares regression. The angle between planes was calculated as the inverse cosine of the dot product of the normalized plane normals.

### Statistics and reproducibility

P values for experiments with multiple conditions were computed using one-way ANOVA. For P values computed using ANOVA, data distribution was assumed to be normal, but this was not formally tested. We used non-parametric two-sided Wilcoxon test or Mann-Whitney test for evaluating P values of paired and unpaired populations respectively. P values were adjusted with Bonferroni correction when performing multiple comparisons. To calculate the 95% confidence interval for each principal components and for the angle between manifolds (**Fig. 6**), we used hierarchical bootstrapping, in which we resampled the population average for each recording location 5000 times. We determined the P value between the different trial types by subtracting 100000 random samples from each distribution and calculating the smallest proportion either above or below 0. Sample sizes were not pre-determined before data acquisition. Sample sizes were balanced between male and female mice, however data for males and females were pooled together since sample sizes were too small to compare across sexes. Data collection and analysis were not performed blind to the conditions of the experiments.

## Supplemental information

**Figure S1.**
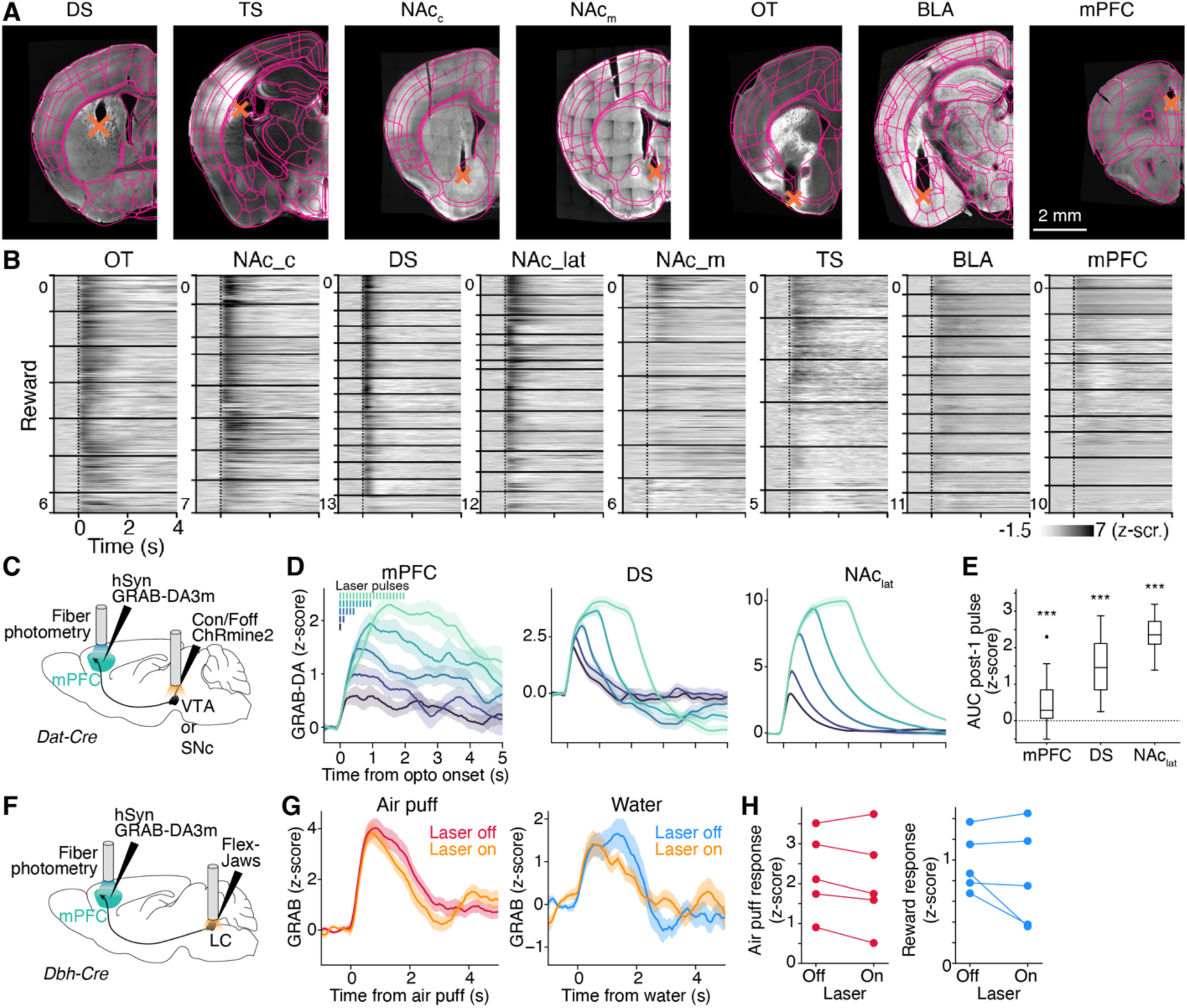
Validation of fiber photometry for measuring dopamine in distinct brain regions, related to Figure 1. **(A)** Examples of fiber placements aligned to the Allen Brain reference atlas (outlined in magenta). Orange X marks indicate the locations of fiber tips. Scale bar: 2 mm. **(B)** Raster plots of single-trial GRAB-DA activity aligned to uncued reward. Horizontal solid lines separate data from different animals. **(C)** Surgical strategy for optogenetic activation of dopamine-expressing neurons in the ventral tegmental area (VTA) or substantia nigra pars compacta (SNc). **(D)** Mean GRAB-DA response to different patterns of light stimulation in DA neurons expressing the opsin ChRmine2.0. **(E)** Mean response to a single 10-ms pulse of light, calculated within a 1-s post-stimulus window. N = 30 trials per condition. ***: P < 0.001 using a Wilcoxon signed-rank test. **(F)** Surgical strategy for optogenetic silencing of noradrenaline-expressing neurons in the locus coeruleus (LC). **(G)** Session average from an example mouse showing GRAB-DA response in the mPFC to air puff or water with (laser on) or without (laser off) LC silencing. **(H)** Mean GRAB-DA response in the mPFC to air puff or water reward with or without LC silencing. N = 5 mice. Data in (C) and (G) are plotted as mean ± s.e.m.

**Figure S2.**
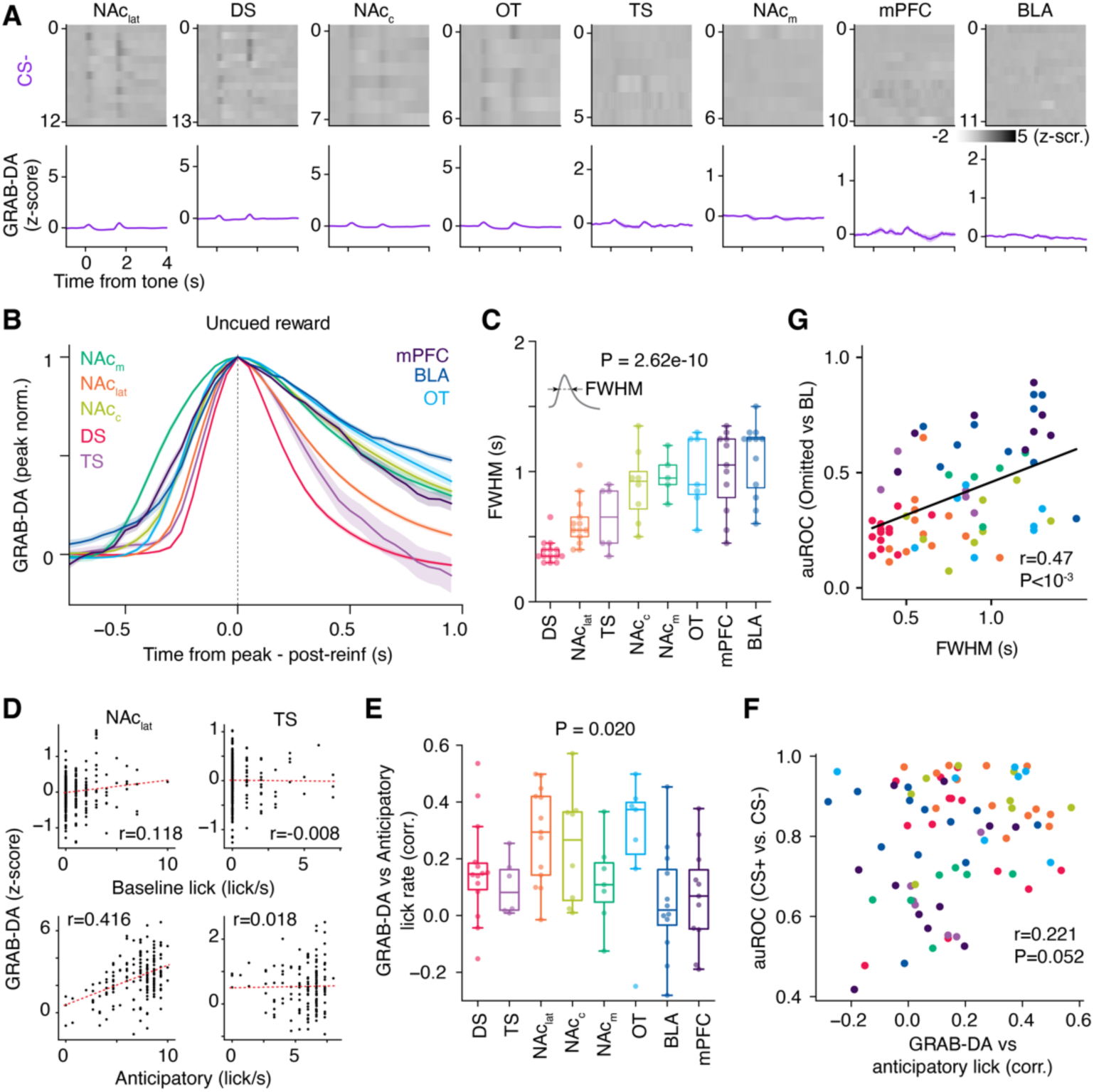
Kinetics of GRAB-DA responses and lick-related activity, related to Figure 2. **(A)** Top: raster plots of GRAB-DA signals across eight brain regions, averaged by animal, during CS- trials. Bottom: population averages for each location. GRAB-DA signals were aligned to tone onset. **(B)** Mean GRAB-DA responses to reward for different recording targets. Responses were peak-normalized and aligned to the time of peak response. **(C)** FWHM (see inset) of the GRAB-DA response to reward across targets. **(D)** GRAB-DA signals as a function of baseline (top) or anticipatory (bottom) lick rate in two example sessions recorded in the NAclat and TS. Pearson *r* values are indicated. **(E)** Correlation between anticipatory lick rate and GRAB-DA signal across recording sites**. (F**) auROC for GRAB-DA responses to CS+ versus CS- plotted against the correlation between GRAB-DA response and anticipatory lick rate. **(G)** Encoding of omitted versus baseline responses plotted as a function of FWHM. Pearson *r* and *p* values are indicated in (F) and (G). Dots in (F) and (G) are color-coded as in (C). *p* values in (C) and (E) were calculated using ANOVA. Data in (A) and (B) are plotted as mean ± s.e.m. N = 78 recording sites.

**Figure S3.**
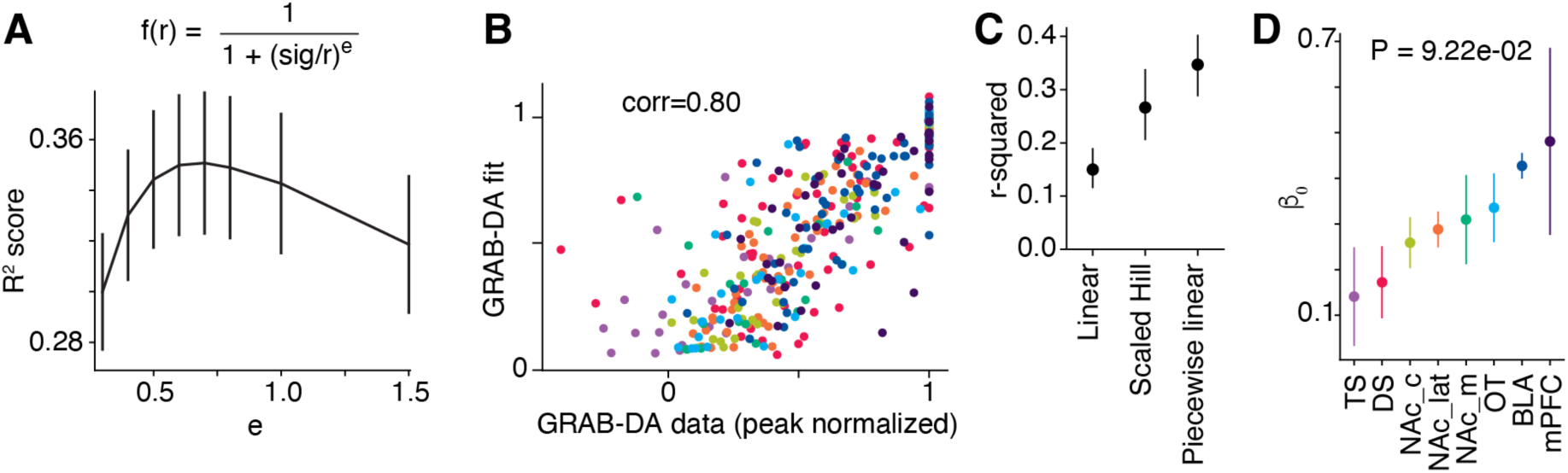
Predicting GRAB-DA responses to different reward sizes, related to Figure 3. **(A)** R-squared values for GRAB-DA responses to uncued rewards predicted by a Hill function *f*(*r*) with Hill coefficient *e*. Data are presented as mean ± s.e.m. **(B)** Correlation between GRAB-DA reward responses predicted by the Hill function and the actual responses. **(C)** R-squared values for different model fits predicting GRAB-DA responses as a function of cued reward size. Data are shown as mean ± 95% confidence interval from bootstrap distributions. **(D)** *β₀* represents the predicted response value at 5 μL reward in the piecewise linear regression. Data are shown as mean ± s.e.m. P value calculated using ANOVA. N = 69 recording locations.

**Figure S4.**
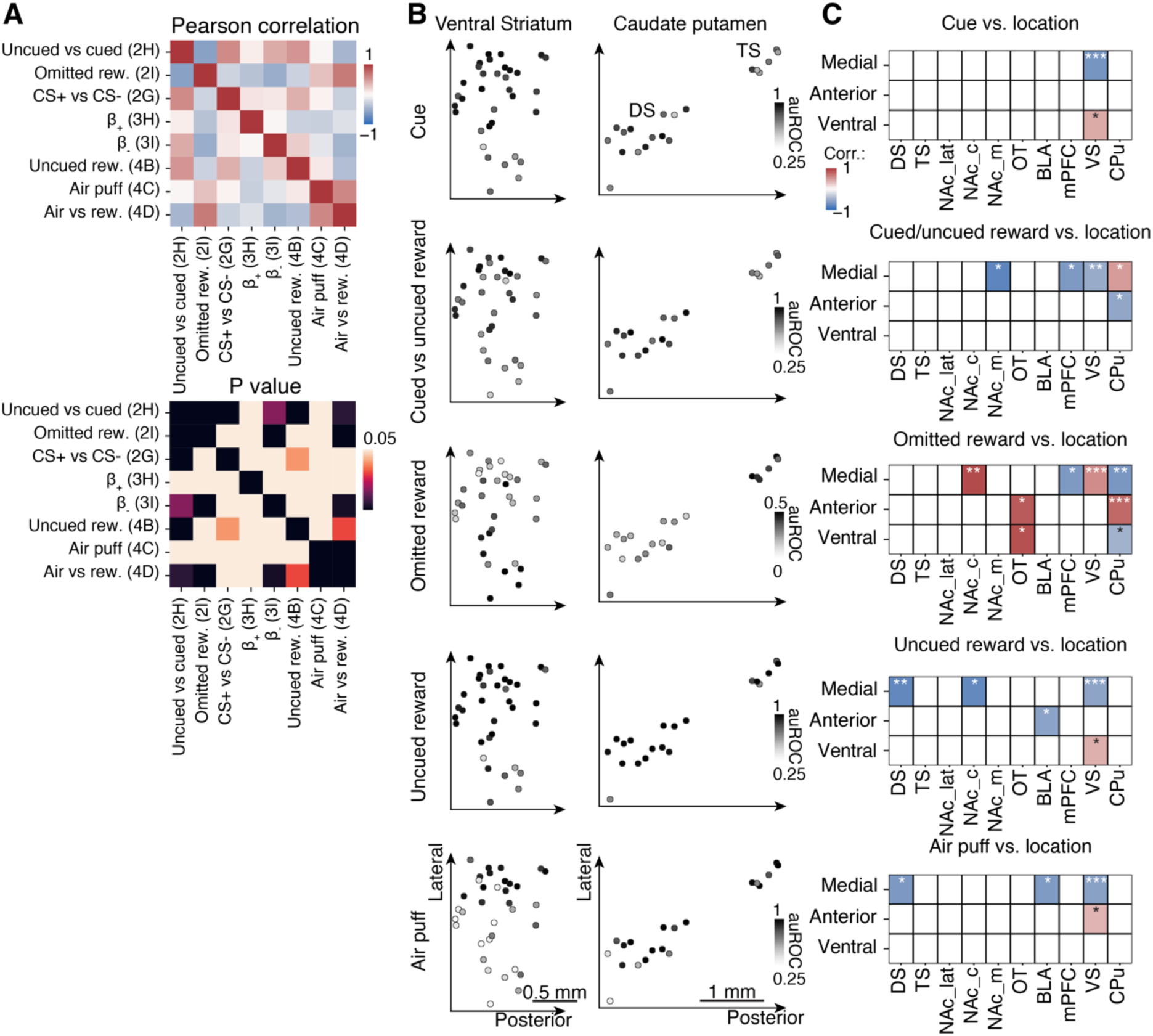
GRAB-DA recording locations and responses during distinct behavioral epochs, related to Figures 2–5. **(A)** Correlation matrix of dopamine response features measured during classical conditioning. Numbers in parentheses refer to corresponding figure panels. The bottom matrix displays corresponding P-values adjusted using Bonferroni correction. **(B)** Recording site locations in the ventral striatum (NAclat, NAcc, NAcm, and OT; N = 35, left) and caudate putamen (DS and TS; N = 20, right), color-coded by auROC of GRAB-DA responses for five behavioral epochs. **(C)** Correlation of auROC values with recording site coordinates along the mediolateral, anteroposterior, and dorsoventral axes. Correlation coefficients were calculated using Pearson correlation. *: P < 0.05; **: P < 0.01; ***: P < 0.001. N = 78 recording sites.

**Figure S5.**
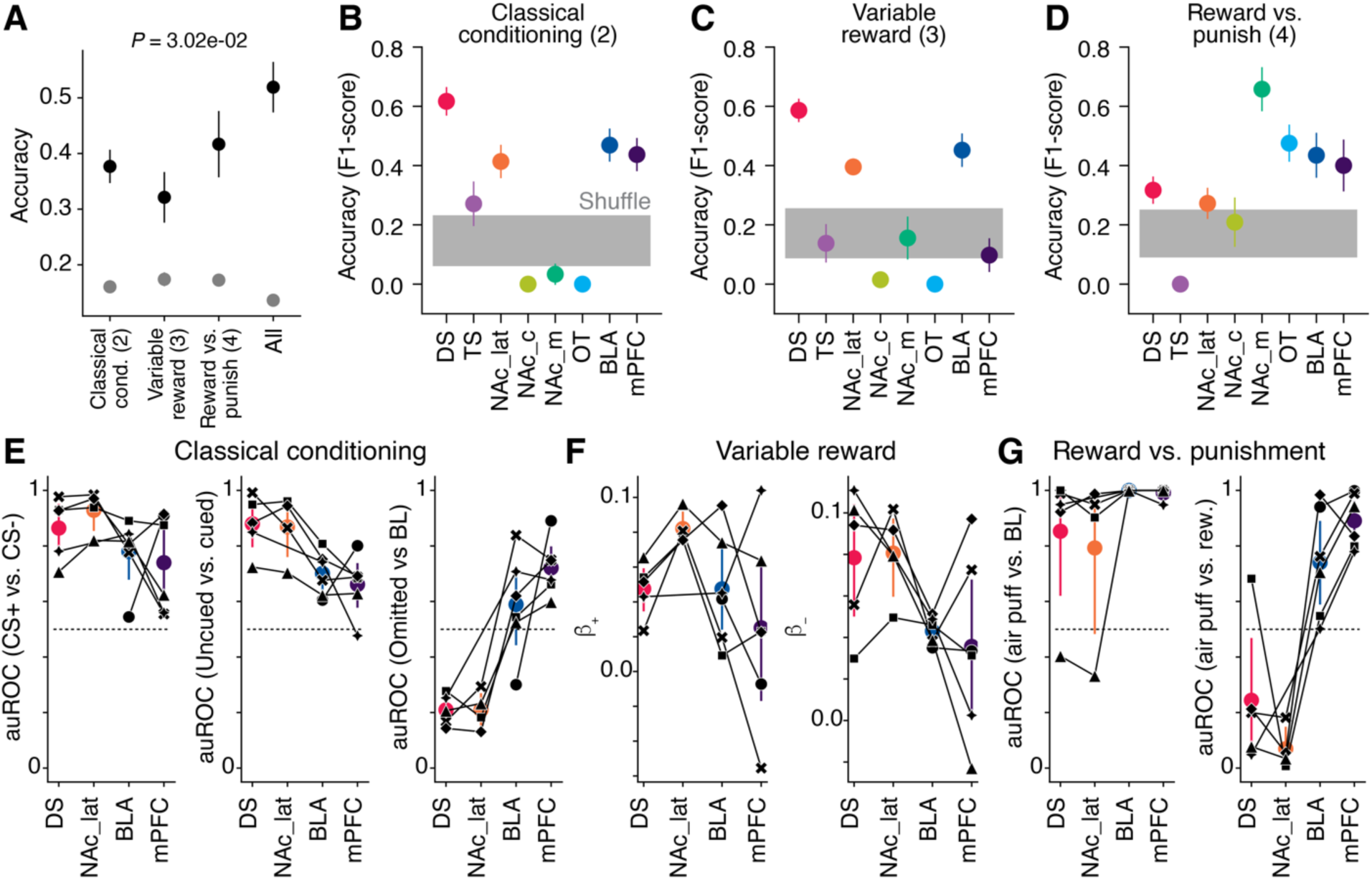
Comparing decoding accuracy of reward features across individual behavioral paradigms, related to Figure 5. (A–D) An SVM classifier was trained to identify recording locations from GRAB-DA response features. **(A)** Mean classification accuracy based on features extracted from the classical conditioning, variable reward, and reward/punishment tasks. P-value calculated with ANOVA comparing classifier performance on real (black) versus shuffled (gray) data. **(B–D)** Classification accuracy for individual target prediction using SVM classifiers trained on GRAB-DA features from each behavioral paradigm (figure # in parentheses). Distribution of cross-validated accuracy scores is shown alongside results from shuffled labels. **(E–G)** GRAB-DA responses as a function of recording location from simultaneous recordings in DS, NAclat, BLA, and mPFC. Colored dots: average per recording site; black lines: individual mouse data. **(E)** auROC for CS− vs. CS+ responses, cued vs. uncued reward, and baseline vs. omitted reward. **(F)** β⁺ and β⁻ represent slopes of the piecewise linear fit for rewards larger and smaller than 5 µL, respectively. **(G)** auROC values comparing responses to air puff vs. baseline and air puff vs. uncued reward. N = 69 recording sites in (A–D) and 7 mice in (E–G). Data are plotted as mean ± s.e.m.

**Figure S6.**
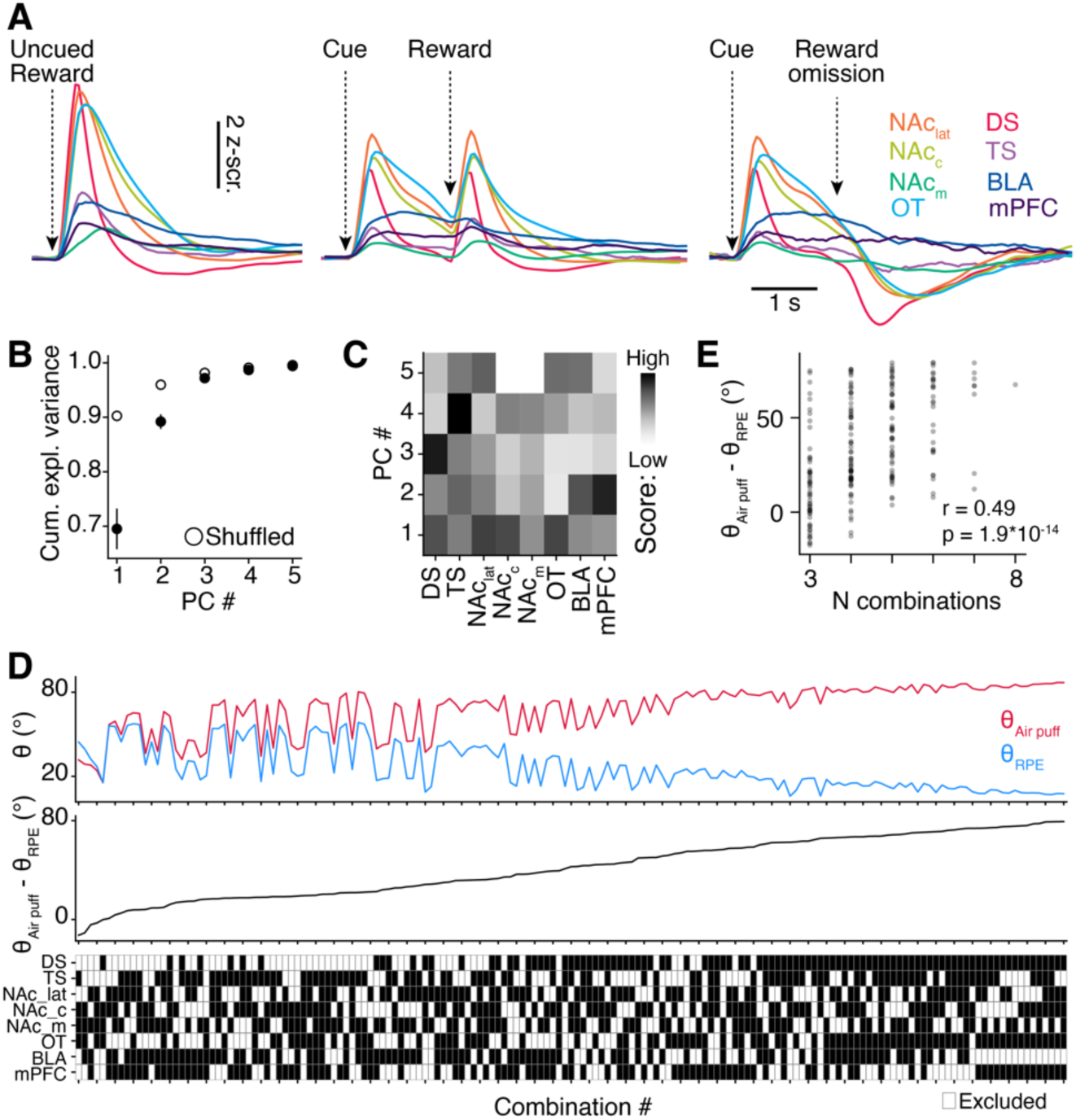
Low-dimensional representation of reinforcement in heterogeneous dopamine signals, related to Figure 6. **(A)** Population average GRAB-DA responses during uncued reward, cued reward, and omitted reward trials. **(B)** Cumulative explained variance as a function of the number of principal components. Data are plotted as mean ± standard deviation. **(C)** Principal components #1-5 used for low-dimensional representation of dopamine signals. **(D)** Top: Angle for pairwise comparison within reward prediction error *θ_RPE_* related signals (blue traces in D) or comparison of air puff and other signals *θ_Air puff_* (red traces in D), as a function of all possible combinations of 4 to 7 recording locations. Middle: Difference between *θ_Air puff_* and *θ_RPE_* as a function of all possible combinations. The larger the angle the more low-dimensional representations can differentiate aversive from reward prediction error events. Bottom: Matrix of all combinations. (**E**) Difference between *θ_Air puff_* and *θ_RPE_* (*Δ*) as a function of number of recording locations. Pearson r: 0.49; p < 10^-14^.

