## Supplementary Figures 1 to 6 for "System-wide dissociation of reward and aversive dopaminergic signals"

### Supplemental information:

Figures S1-S6

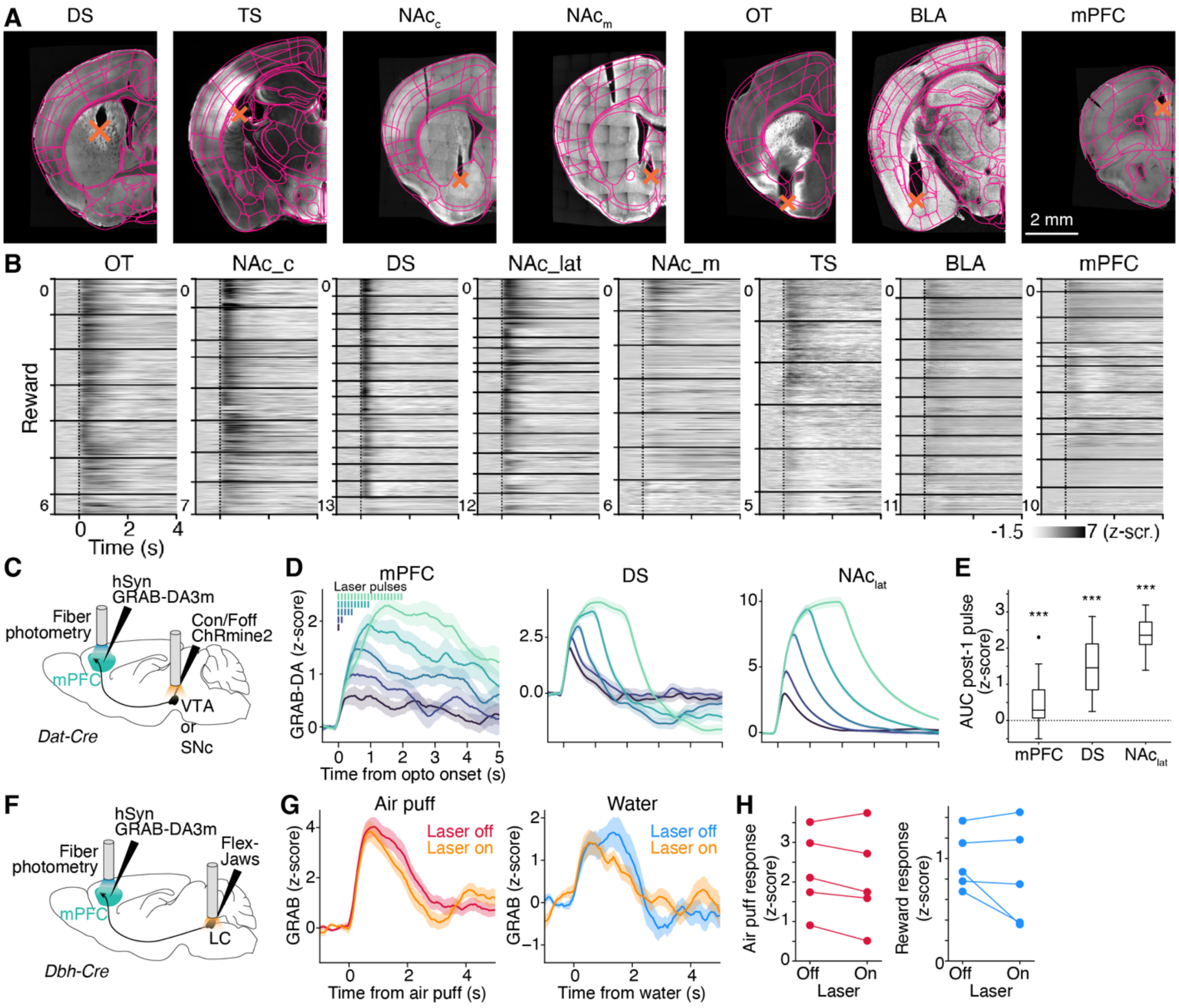


**Figure S1. Validation of fiber photometry for measuring dopamine in distinct brain regions, related to Figure 1. (A)** Examples of fiber placements aligned to the Allen Brain reference atlas (outlined in magenta). Orange X marks indicate the locations of fiber tips. Scale bar: 2 mm. **(B)** Raster plots of single-trial GRAB-DA activity aligned to uncued reward. Horizontal solid lines separate data from different animals. **(C)** Surgical strategy for optogenetic activation of dopamine-expressing neurons in the ventral tegmental area (VTA) or substantia nigra pars compacta (SNc). **(D)** Mean GRAB-DA response to different patterns of light stimulation in DA neurons expressing the opsin ChRmine2.0. **(E)** Mean response to a single 10-ms pulse of light, calculated within a 1-s post-stimulus window. N = 30 trials per condition. ***: $P < 0.001$ using a Wilcoxon signed-rank test. **(F)** Surgical strategy for optogenetic silencing of noradrenaline-expressing neurons in the locus coeruleus (LC). **(G)** Session average from an example mouse showing GRAB-DA response in the mPFC to air puff or water with (laser on) or without (laser off) LC silencing. **(H)** Mean GRAB-DA response in the mPFC to air puff or water reward with or without LC silencing. N = 5 mice. Data in (C) and (G) are plotted as mean ± s.e.m.

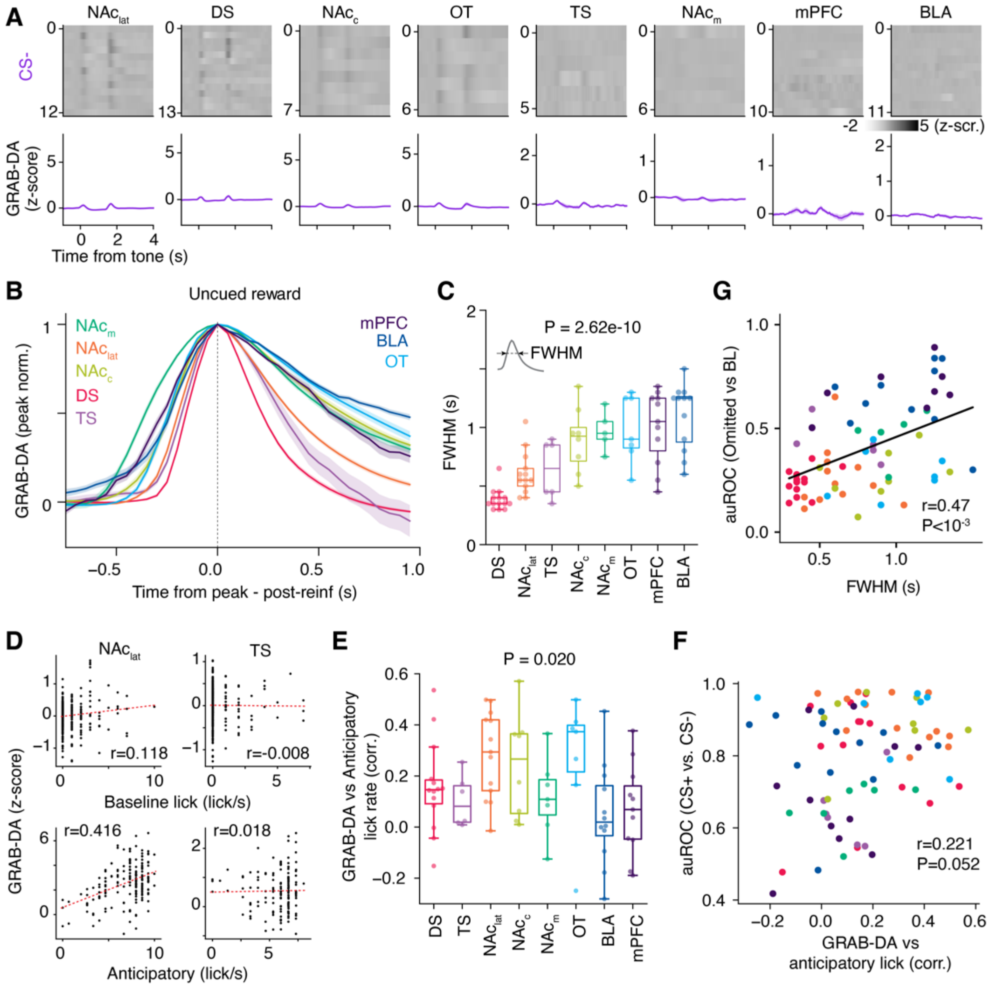


**Figure S2. Kinetics of GRAB-DA responses and lick-related activity, related to Figure 2. (A)** Top: raster plots of GRAB-DA signals across eight brain regions, averaged by animal, during CS- trials. Bottom: population averages for each location. GRAB-DA signals were aligned to tone onset. **(B)** Mean GRAB-DA responses to reward for different recording targets. Responses were peak-normalized and aligned to the time of peak response. **(C)** FWHM (see inset) of the GRAB-DA response to reward across targets. **(D)** GRAB-DA signals as a function of baseline (top) or anticipatory (bottom) lick rate in two example sessions recorded in the NAclat and TS. Pearson *r* values are indicated. **(E)** Correlation between anticipatory lick rate and GRAB-DA signal across recording sites**. (F)** auROC for GRAB-DA responses to CS+ versus CS- plotted against the correlation between GRAB-DA response and anticipatory lick rate. **(G)** Encoding of omitted versus baseline responses plotted as a function of FWHM. Pearson *r* and *p* values are indicated in (F) and (G). Dots in (F) and (G) are color-coded as in (C). *p* values in (C) and (E) were calculated using ANOVA. Data in (A) and (B) are plotted as mean ± s.e.m. N = 78 recording sites.

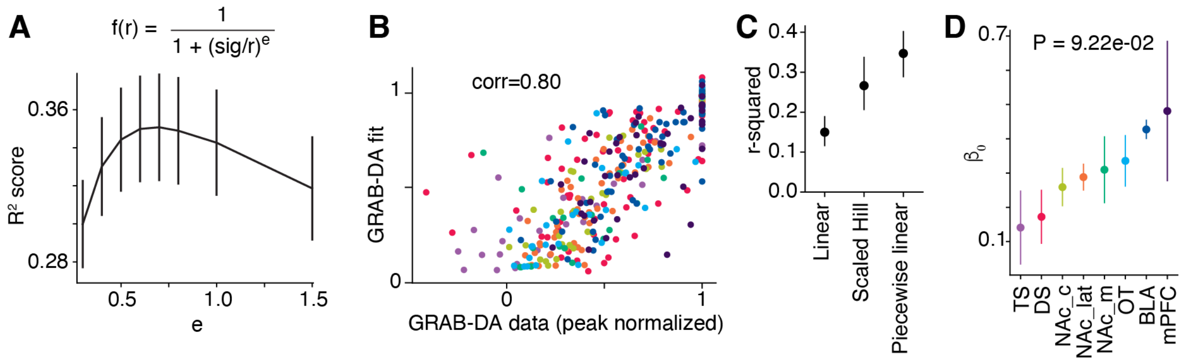


**Figure S3. Predicting GRAB-DA responses to different reward sizes, related to Figure 3. (A)** R-squared values for GRAB-DA responses to uncued rewards predicted by a Hill function $f(r)$ with Hill coefficient $e$. Data are presented as mean ± s.e.m. **(B)** Correlation between GRAB-DA reward responses predicted by the Hill function and the actual responses. **(C)** R-squared values for different model fits predicting GRAB-DA responses as a function of cued reward size. Data are shown as mean ± 95% confidence interval from bootstrap distributions. **(D)** $\beta_0$ represents the predicted response value at 5 µL reward in the piecewise linear regression. Data are shown as mean ± s.e.m. P value calculated using ANOVA. N = 69 recording locations.

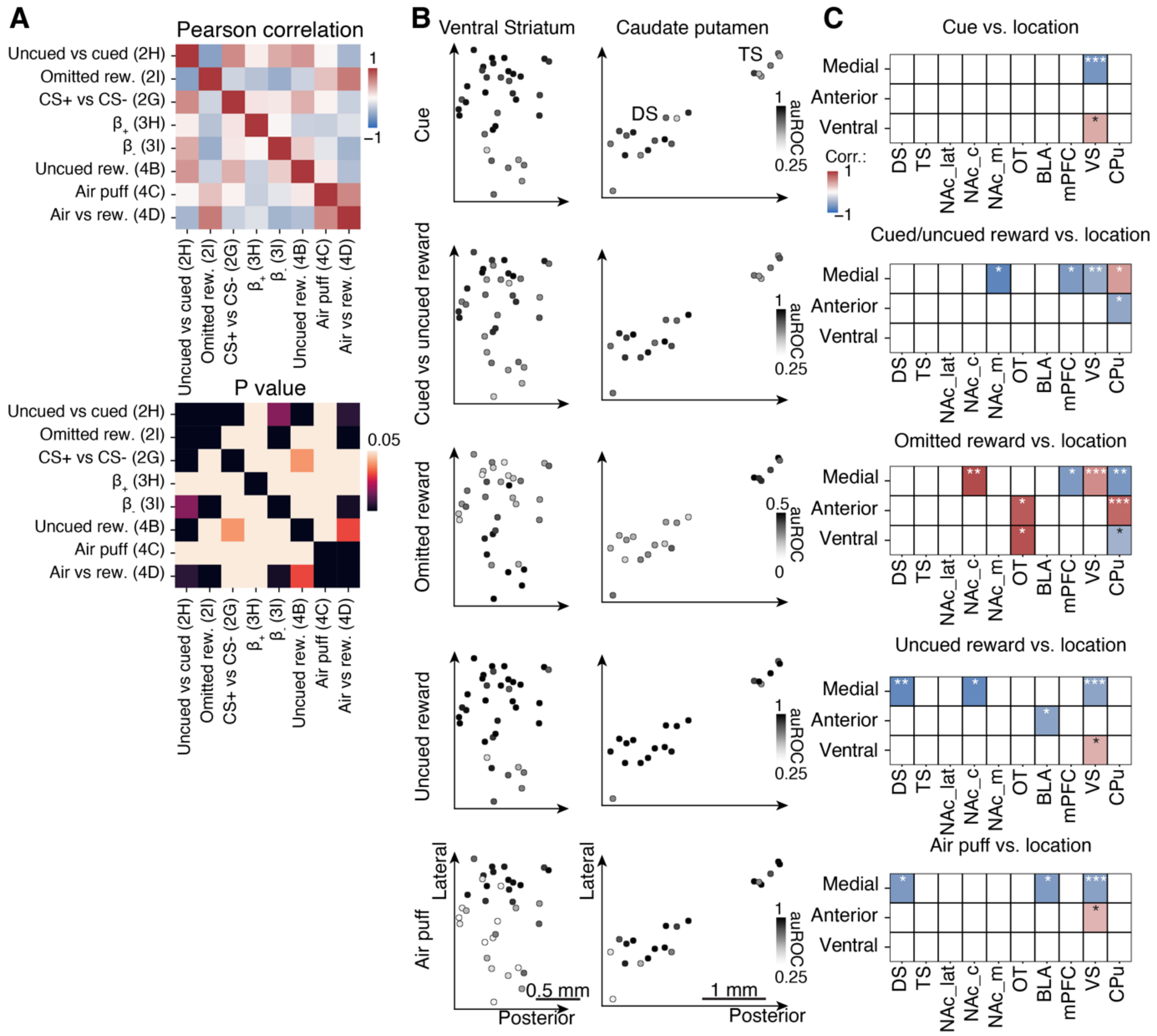


**Figure S4. GRAB-DA recording locations and responses during distinct behavioral epochs, related to Figures 2–5. (A)** Correlation matrix of dopamine response features measured during classical conditioning. Numbers in parentheses refer to corresponding figure panels. The bottom matrix displays corresponding P-values adjusted using Bonferroni correction. **(B)** Recording site locations in the ventral striatum (NAclat, NAcc, NAcm, and OT; N = 35, left) and caudate putamen (DS and TS; N = 20, right), color-coded by auROC of GRAB-DA responses for five behavioral epochs. **(C)** Correlation of auROC values with recording site coordinates along the mediolateral, anteroposterior, and dorsoventral axes. Correlation coefficients were calculated using Pearson correlation. *: $P < 0.05$; **: $P < 0.01$; ***: $P < 0.001$. N = 78 recording sites.

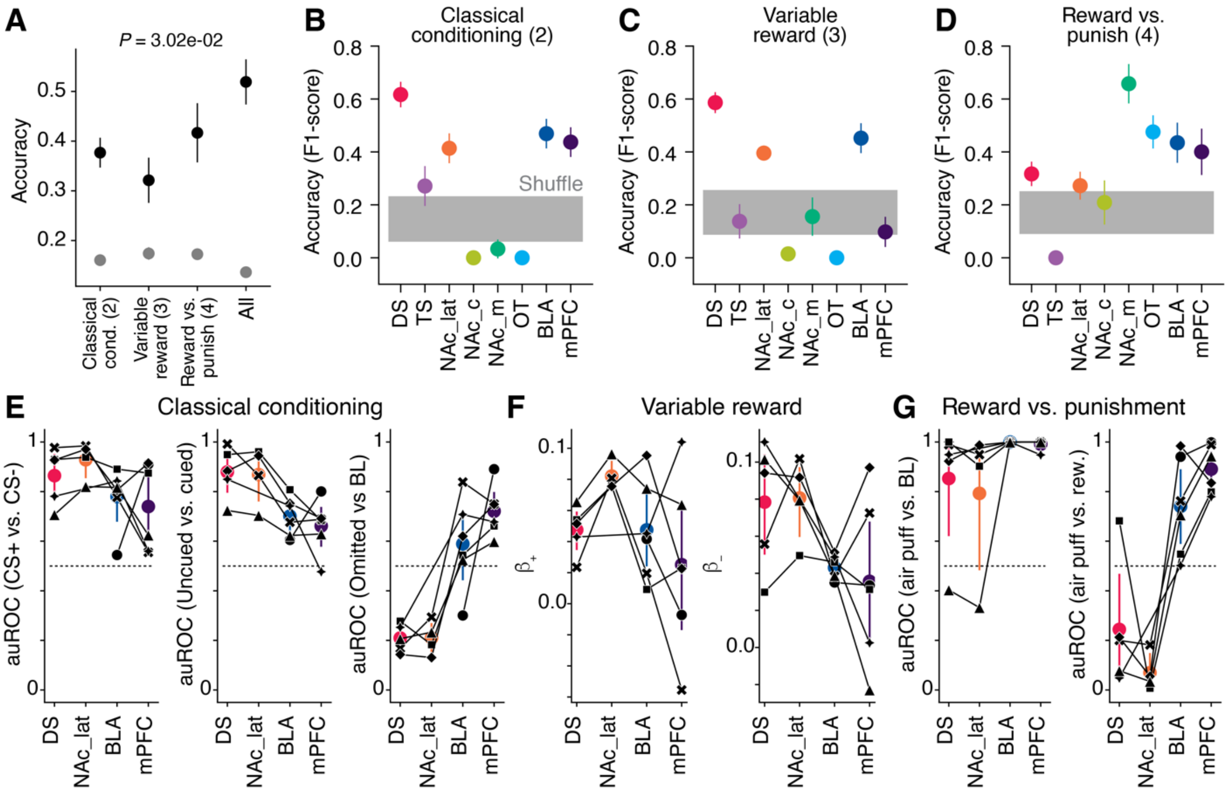


**Figure S5. Comparing decoding accuracy of reward features across individual behavioral paradigms, related to Figure 5.** **(A–D)** An SVM classifier was trained to identify recording locations from GRAB-DA response features. **(A)** Mean classification accuracy based on features extracted from the classical conditioning, variable reward, and reward/punishment tasks. P-value calculated with ANOVA comparing classifier performance on real (black) versus shuffled (gray) data. **(B–D)** Classification accuracy for individual target prediction using SVM classifiers trained on GRAB-DA features from each behavioral paradigm (figure # in parentheses). Distribution of cross-validated accuracy scores is shown alongside results from shuffled labels. **(E–G)** GRAB-DA responses as a function of recording location from simultaneous recordings in DS, NAclat, BLA, and mPFC. Colored dots: average per recording site; black lines: individual mouse data. **(E)** auROC for CS− vs. CS+ responses, cued vs. uncued reward, and baseline vs. omitted reward. **(F)** $\beta^{+}$ and $\beta^{-}$ represent slopes of the piecewise linear fit for rewards larger and smaller than 5 µL, respectively. **(G)** auROC values comparing responses to air puff vs. baseline and air puff vs. uncued reward. N = 69 recording sites in (A–D) and 7 mice in (E–G). Data are plotted as mean ± s.e.m.

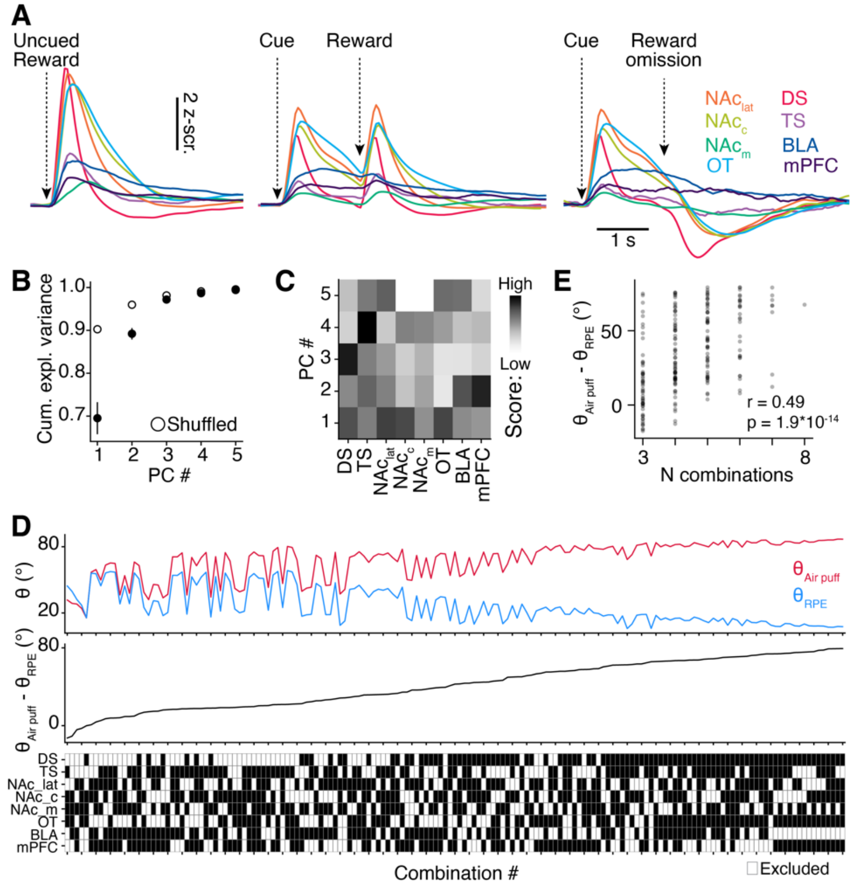


**Figure S6. Low-dimensional representation of reinforcement in heterogeneous dopamine signals, related to Figure 6.** **(A)** Population average GRAB-DA responses during uncued reward, cued reward, and omitted reward trials. **(B)** Cumulative explained variance as a function of the number of principal components. Data are plotted as mean ± standard deviation. **(C)** Principal components #1-5 used for low-dimensional representation of dopamine signals. **(D)** Top: Angle for pairwise comparison within reward prediction error $\theta_{RPE}$ related signals (blue traces in D) or comparison of air puff and other signals $\theta_{Air\ puff}$ (red traces in D), as a function of all possible combinations of 4 to 7 recording locations. Middle: Difference between $\theta_{Air\ puff}$ and $\theta_{RPE}$ as a function of all possible combinations. The larger the angle the more low-dimensional representations can differentiate aversive from reward prediction error events. Bottom: Matrix of all combinations. (**E**) Difference between $\theta_{Air\ puff}$ and $\theta_{RPE}$ ($\Delta$) as a function of number of recording locations. Pearson r: 0.49; $p < 10^{-14}$.
